# The *Mycobacterium tuberculosis* ESX-5 secretion system enables carbon source utilization and growth in mice

**DOI:** 10.1101/2025.10.28.685107

**Authors:** Alisha M. Block, Rashmi Ravindran Nair, Virginia Meikle, Parker C. Wiegert, Dylan W. White, Leanne Zhang, Michael Niederweis, Anna D. Tischler

## Abstract

*Mycobacterium tuberculosis* uses several ESX type VII protein secretion systems for pathogenesis. *M. tuberculosis* ESX-5 is poorly characterized because it is essential for growth in standard lab culture conditions. To circumvent ESX-5 essentiality, we made an *M. tuberculosis* strain in which the central ESX-5 membrane component EccD_5_ can be conditionally depleted. Here, we use this strain to demonstrate that *M. tuberculosis* requires the ESX-5 secretion system to grow using specific carbon sources *in vitro*, to grow in cultured macrophages, and to replicate and disseminate in aerosol-infected mice. *M. tuberculosis* requires ESX-5 to use glycerol or glucose as the sole carbon source. Use of glycerol and glucose also depends on the outer membrane protein PPE51. We show that *M. tuberculosis* requires ESX-5 activity for outer membrane export and surface exposure of PPE51. Expression of the outer membrane porin MspA enabled growth of ESX-5 deficient *M. tuberculosis* on glycerol, suggesting that the main function of ESX-5 *in vitro* is to export nutrient transporters to the outer membrane. Importantly, depletion of EccD_5_ in acutely infected mice resulted in clearance of *M. tuberculosis* from lung tissues, demonstrating the critical importance of ESX-5 activity during infection. Our findings suggest that ESX-5 promotes *M. tuberculosis* pathogenesis by mediating export of outer membrane proteins that enable nutrient acquisition.

**Importance:** *Mycobacterium tuberculosis* ESX type VII secretion systems play important roles in pathogenesis, but the functions of ESX-5 are not well characterized because it is essential for growth in standard lab culture conditions. We used a strain that conditionally expresses a central membrane component of the ESX-5 secretion apparatus to determine how ESX-5 impacts growth in lab cultures and in a mouse infection model. We found that *M. tuberculosis* requires ESX-5 to grow using several carbon sources and to grow in the lungs of infected mice. Inhibiting production of the ESX-5 secretion system in mice also led to clearance of *M. tuberculosis* from lung tissues. Our results demonstrate that the *M. tuberculosis* ESX-5 system is a critical virulence factor and suggest that ESX-5 is a strong candidate for anti-tubercular drug development.

## Introduction

*Mycobacterium tuberculosis* encodes five ESX type VII protein secretion systems that play critical roles in pathogenesis by exporting proteins that manipulate the functions of infected phagocytes (1, 2). ESX-1, ESX-2 and ESX-4 all participate in permeabilizing the phagosomal membrane, enabling *M. tuberculosis* and secreted molecules to access the host cell cytosol (3, 4). ESX-1 induces pyroptosis of infected macrophages by disrupting the plasma membrane, which activates the NLRP3 inflammasome (5-7). ESX-4 exports the tuberculosis necrotizing toxin (CpnT or TNT) that triggers macrophage necrosis (3, 8). *M. tuberculosis* requires ESX-3 to acquire iron (9, 10) and to prevent ESCRT-dependent phagosomal membrane repair, which inhibits antigen processing and T cell activation (11, 12). The related pathogen *Mycobacterium marinum* requires both ESX-5 and ESX-1 to induce inflammasome activation and pyroptotic death of infected macrophages (13). However, analysis of ESX-5 function in *M. tuberculosis* pathogenesis has been complicated by the fact that ESX-5 is essential for *in vitro* growth.

Genetic evidence suggests that *M. tuberculosis* requires ESX-5 for growth in standard lab culture conditions. Genes encoding ESX-5 conserved components that form an inner membrane secretion complex [EccB_5_, EccC_5_, EccD_5_, EccE_5_ and MycP_5_ (14, 15)] are essential for growth of *M. tuberculosis* lab strains and clinical isolates in standard culture media based on genome-wide Tn-seq screens (16, 17). The *eccB_5_*, *eccC_5_* and *eccD_5_* genes were confirmed to be essential as these genes could not be deleted without a complementing copy provided *in trans* (18, 19). Both *eccC_5_* and *mycP_5_* are similarly essential in *Mycobacterium marinum* (20). *M. tuberculosis* Δ*eccD_5_* and Δ*esx-5* mutants, in which the entire *esx-5* locus is deleted, were described (21-23). However, one Δ*esx-5* strain has a secondary mutation that prevents production of phthiocerol dimycocerosate (PDIM) (23), an outer membrane lipid essential for virulence (24, 25). Rigorous genetic analyses were not done to confirm the absence of compensatory mutations in the other Δ*esx-5* mutants or the Δ*eccD_5_* mutant (21, 22). The Δ*eccD_5_* mutant failed to replicate in intravenously infected severe combined immunodeficiency (SCID) mice, but this phenotype was not fully complemented (21). A Δ*esx-5* mutant similarly failed to replicate in intravenously infected SCID or *Rag*^-/-^ mice (22), but complementation was not done. Thus, ESX-5 likely plays a critical role in *M. tuberculosis* physiology that promotes growth in the host, but this has yet to be tested directly in an immune-competent animal model.

Clues to the function of ESX-5 in mycobacterial physiology were provided by studies in *M. marinum*, which suggested that ESX-5 enables *in vitro* growth by increasing permeability of the mycobacterial outer membrane to nutrients (20). The *eccC_5_* or *mycP_5_* genes could be deleted in *M. marinum* expressing the *M. smegmatis* porin MspA (20), which transports small hydrophilic nutrients including glucose and serine across the *M. smegmatis* outer membrane (26, 27). In *M. marinum*, *eccC_5_* could also be disrupted in strains lacking PDIM (20). The PDIM lipid decreases permeability of the mycobacterial outer membrane to small molecules, including nutrients and antibiotics (28-31). Taken together, these data imply that ESX-5 normally functions to export proteins that enhance nutrient transport through the *M. marinum* outer membrane.

Based on studies done in *M. marinum*, ESX-5 is predicted to export most proteins of the mycobacteria-specific PE and PPE families (20, 32, 33). The *M. tuberculosis* H37Rv reference genome encodes 99 PE proteins and 69 PPE proteins, representing ∼8% of the genome coding capacity (34, 35). Most of these *M. tuberculosis* PE and PPE proteins are likely to be exported via ESX-5, as immune recognition of conserved PE and PPE epitopes requires a functional ESX-5 secretion system (36). A subset of PE and PPE proteins were implicated in uptake of small molecule nutrients including heme, Ca^2+^, glucose, glycerol and various disaccharides (28, 37-43). Some PPE proteins linked to nutrient uptake localize to the *M. tuberculosis* cell surface where they have been proposed to act either as receptors that capture nutrients or as porin-like channels that facilitate nutrient transport through the outer membrane (28, 37, 41). As ESX-5 is predicted to export most PE and PPE proteins, a major function of ESX-5 that promotes *M. tuberculosis* growth *in vitro* may be to export these proteins for nutrient acquisition. However, the molecular mechanisms by which ESX-5 promotes *in vitro* growth and the role of ESX-5 in *M. tuberculosis* pathogenesis have not been determined.

Here we use a *M. tuberculosis* strain that conditionally expresses the ESX-5 core component EccD_5_ to determine the roles of ESX-5 in growth *in vitro*, in macrophages and in mice. We find that EccD_5_ depletion prevents growth on glucose and glycerol, the primary carbon sources in standard lab media. We demonstrate that *M. tuberculosis* requires ESX-5 for outer membrane localization of PPE51, which was previously implicated in glucose and glycerol uptake (28), suggesting that ESX-5 exports PPE51 to enable growth on glucose and glycerol. In macrophages, we find that *M. tuberculosis* requires ESX-5 for replication and for induction of inflammatory cell death. Finally, in mice, we find that EccD_5_ depletion prevents *M. tuberculosis* replication in the lungs and dissemination to the spleen and causes clearance of the bacteria during acute infection. Collectively, our findings suggest that ESX-5 promotes *M. tuberculosis* pathogenesis in part by exporting outer membrane proteins that enable nutrient acquisition.

## Results

To circumvent essentiality of ESX-5 for *M. tuberculosis* growth *in vitro*, we generated an *eccD_5_* Tet-OFF strain in which the native *eccD_5_* gene is deleted and *eccD_5_* is expressed *in trans* from a tetracycline-repressible (Tet-OFF) promoter (19). We showed that *eccD_5_* is transcriptionally repressed, EccD_5_ protein is depleted, and secretion of two known ESX-5 substrates EsxN and PPE41 is reduced upon treatment of *eccD_5_* Tet-OFF with anhydrotetracycline (Atc) (3, 19). We verified by whole genome sequencing that our *eccD_5_* Tet-OFF strain does not harbor mutations in genes required to produce the PDIM lipid. We also confirmed by analysis of ^14^C-propionate labeled lipid extracts that *eccD_5_* Tet-OFF produces PDIM (**Fig S1A**). We now exploit this *eccD_5_* Tet-OFF strain to determine the role of the ESX-5 secretion system in *M. tuberculosis* growth *in vitro* and in mice.

### *M. tuberculosis* requires ESX-5 to grow on specific carbon sources *in vitro*

Previous studies in *M. marium* implicated ESX-5 in growth on esterified fatty acids (20). *M. marinum* also likely requires ESX-5 for growth on small hydrophilic carbon sources, as ESX-5 core components could be deleted in a strain expressing the *M. smegmatis* porin MspA (20, 26, 27). To determine the role of *M. tuberculosis* ESX-5 in carbon source utilization, we analyzed growth of our *eccD_5_* Tet-OFF strain in Middlebrook 7H9 base medium supplemented with various sole carbon sources. The 7H9 base does not support growth of WT *M. tuberculosis* Erdman or *eccD_5_* Tet-OFF (**Fig 1A**). WT Erdman and *eccD_5_* Tet-OFF grew on all carbon sources tested (**Fig 1B-E**), though growth of the *eccD_5_* Tet-OFF strain was significantly delayed on glycerol, even without Atc (**Fig 1B**). Depletion of EccD_5_ with Atc prevented *M. tuberculosis* growth on glycerol (**Fig 1B**), glucose (**Fig 1C**) and the esterified fatty acid Tween-40 (Tw-40, **Fig 1D**). We observed similar growth defects on glycerol or glucose in a minimal salts medium when EccD_5_ was depleted (**Fig S2**). In contrast, EccD_5_ depletion only slightly reduced the growth rate on cholesterol as the sole carbon source (**Fig 1E**), suggesting that *M. tuberculosis* does not require ESX-5 to use cholesterol or other nutrients in Middlebrook 7H9. We confirmed that EccD_5_ was efficiently depleted from the *eccD_5_* Tet-OFF strain grown in 7H9 with cholesterol + Atc (**Fig 1F**). These data indicate that *M. tuberculosis* requires ESX-5 to grow using glycerol, glucose or Tw-40 as the sole carbon source. As glycerol and glucose are the primary carbon sources in Middlebrook media, our data suggest that ESX-5 is essential *in vitro* because it is required to use these carbon sources.

**Figure 1.**
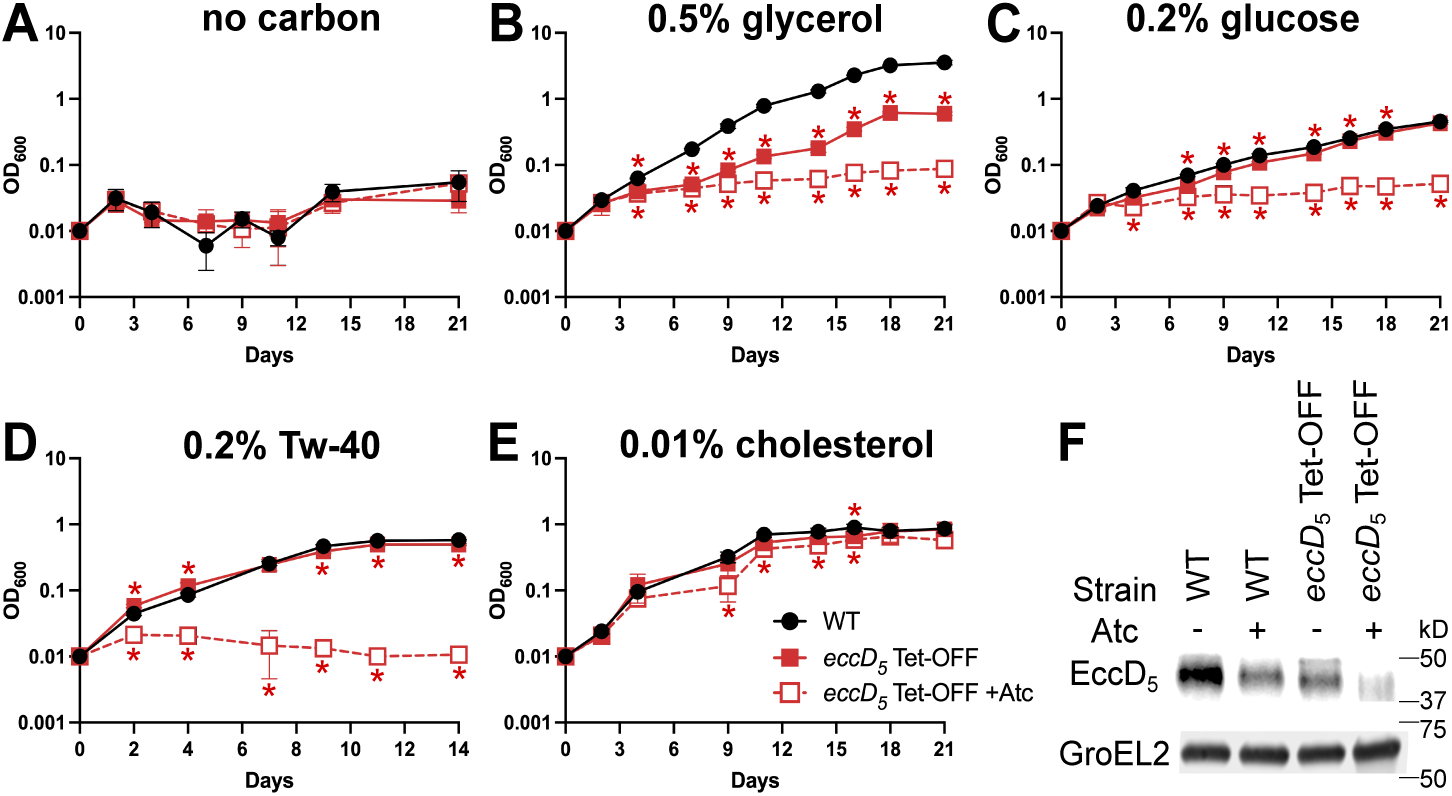
*M. tuberculosis* requires ESX-5 for *in vitro* growth on various carbon sources. **(A-E)** WT Erdman and *eccD_5_* Tet-OFF were grown in complete Middlebrook 7H9 ± 100 ng/ml Atc to mid-exponential phase, then washed and diluted to OD_600_ = 0.01 in triplicate in Middlebrook 7H9 base with 0.01% tyloxapol and no added carbon (**A**), 0.5% glycerol (**B**), 0.2% glucose (**C**), 0.2% Tween-40 (**D**), or 0.01% cholesterol (**E**) ± 100 ng/ml anhydrotetracycline (Atc). In **E**, fresh cholesterol was added at 0.01% every 2-3 days. Growth was monitored by OD_600_ measurements every 2-3 days. Fresh Atc (100 ng/ml) was added to +Atc cultures every 7 days. Data are means ± standard deviations. A one-way ANOVA with Dunnett’s corrections was used to compare *eccD_5_* Tet-OFF ± Atc and to WT (\**p*<0.05). (**F**) Proteins were extracted from cholesterol cultures at day 21. EccD_5_ and GroEL2 were detected in whole cell lysates (11.6 μg total protein) by Western blotting.

### *M. tuberculosis* requires ESX-5 for nutrient uptake through the mycobacterial outer membrane

Loss of ESX-5 function in *M. marinum* could be complemented by expression of the MspA outer membrane porin from *M. smegmatis* (20). We similarly attempted to complement *in vitro* growth of our *eccD_5_* Tet-OFF strain with MspA. Expression of MspA *in trans* from pMV- *mspA* did not affect growth of WT *M. tuberculosis* (**Fig 2A**), but significantly improved growth of EccD_5_-depleted *M. tuberculosis* in 7H9 with glycerol as the sole carbon source (**Fig 2C**). Western blotting confirmed that EccD_5_ was depleted by Atc in the *eccD_5_* Tet-OFF pMV-*mspA* strain (**Fig S3A**). The *eccD_5_* Tet-OFF pMV261 and *eccD_5_* Tet-OFF pMV-*mspA* strains both produce PDIM (**Fig S1B**), so growth of the MspA-expressing strain on glycerol is not due to loss of the PDIM lipid. However, even without Atc we observed an extended lag phase for the *eccD_5_* Tet-OFF pMV-*mspA* strain relative to the *eccD_5_* Tet-OFF pMV261 and WT controls (**Fig 2C**).

**Figure 2.**
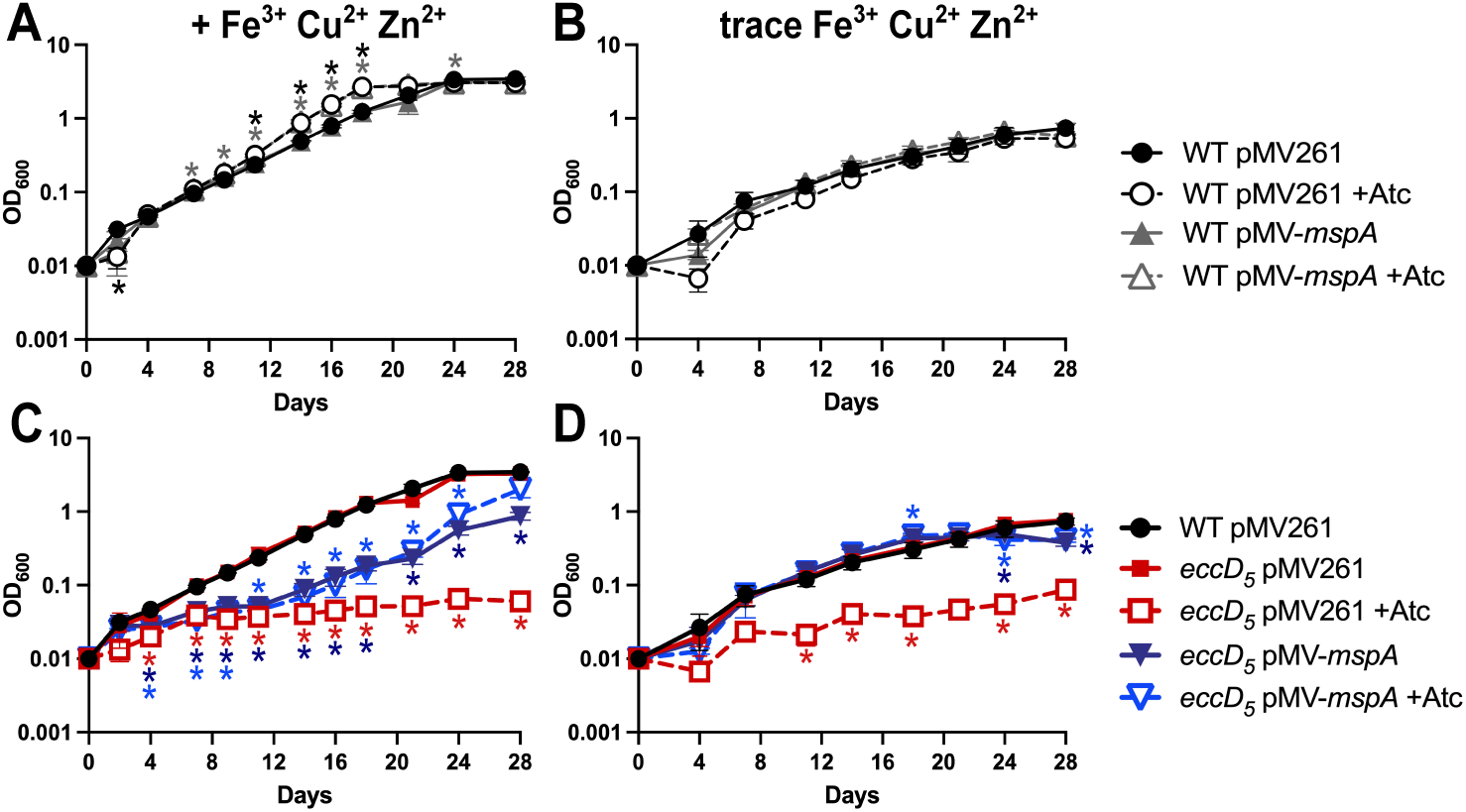
Expression of the *M. smegmatis* porin MspA rescues growth of EccD_5_-deficient *M. tuberculosis* on glycerol. The indicated strains were grown in complete Middlebrook 7H9 ± 100 ng/ml Atc to mid-exponential phase, then washed and diluted in triplicate to OD_600_ = 0.01 in Middlebrook 7H9 base with 0.01% tyloxapol and 0.5% glycerol (**A, C**) or in home-made Middlebrook 7H9 with trace Fe^3+^, Cu^2+^ and Zn^2+^ with 0.01% tyloxapol and 0.5% glycerol (**B, D**) ± 100 ng/ml anhydrotetracycline (Atc). Growth was monitored by OD_600_ measurements every 2-3 days. Fresh Atc (100 ng/ml) was added to +Atc cultures every 7 days. Data are means ± standard deviations. Statistical analysis was done to compare each strain and condition to the WT pMV261 untreated control (**A-B**) or to compare each strain and condition to the *eccD_5_* Tet-OFF pMV261 untreated control (**C-D**) (\**p*<0.05; two-way ANOVA with a simple effects model and Dunnett’s correction).

In addition to transporting small molecule nutrients, MspA also increases uptake of the heavy metals Fe^3+^ and Cu^2+^ (44, 45). Excess Cu^2+^ and Zn^2+^ can be toxic to *M. tuberculosis* (45, 46). To test if metal toxicity contributes to reduced growth of the *eccD_5_* Tet-OFF pMV-*mspA* strain, we conducted growth curves in self-made 7H9 medium without added Fe^3+^, Cu^2+^, or Zn^2+^, which we expect contains trace amounts of these metals (10, 47, 48). In 7H9 with trace metals and glycerol as the sole carbon source, the WT controls grew slowly and reached a lower cell density (**Fig 2B**), but the *eccD_5_* Tet-OFF pMV-*mspA* strain grew similarly to WT and *eccD_5_* Tet-OFF pMV261 controls (**Figs 2D**). Importantly, growth of the *eccD_5_* Tet-OFF pMV-*mspA* strain was not inhibited by Atc (**Fig 2D**). In contrast, growth of the *eccD_5_* Tet-OFF pMV261 empty vector control was inhibited by Atc (**Fig 2D**). Western blotting controls showed that EccD_5_ was depleted by Atc in these experiments (**Figs S3B**). Growth curves in 7H9 with trace metals and no added carbon showed that MspA does not enable use of other carbon sources in the 7H9 base medium (**Fig S3C-D**). Collectively, these data suggest that *M. tuberculosis* requires EccD_5_ for uptake of glycerol across the mycobacterial outer membrane. These data also suggest that *M. tuberculosis* requires ESX-5 to detoxify excess heavy metals, which may include Fe^3+^, Cu^2+^, and/or Zn^2+^.

### *M. tuberculosis* requires ESX-5 for outer membrane localization of PPE51

*M. tuberculosis* requires the PPE51 protein, which localizes to the cell surface, for uptake of and growth on glycerol and glucose (28). As we observed growth defects of EccD_5_-depleted *M. tuberculosis* on these carbon sources, we predicted that ESX-5 would export PPE51 to the outer membrane. To test this, we expressed PPE51 with a C-terminal 6xHis tag (PPE51_His6_) in WT Erdman and *eccD_5_* Tet-OFF and performed Western blotting of soluble and membrane fractions. PPE51_His6_ was detected in the membrane fraction of WT *M. tuberculosis* and the *eccD_5_* Tet-OFF strain grown without Atc but was undetectable in the membrane fraction when EccD_5_ was depleted with Atc (**Fig 3A**). PPE51_His6_ was also not detected in whole cell lysates upon EccD_5_ depletion (**Fig 3A**), suggesting that PPE51_His6_ is not stable if it cannot be exported by ESX-5. The GlcB soluble and LpqH membrane fraction controls confirmed efficient fractionation and equivalent loading (**Fig 3A**). These data suggest that *M. tuberculosis* requires ESX-5 to export PPE51 to the membrane fraction.

**Figure 3.**
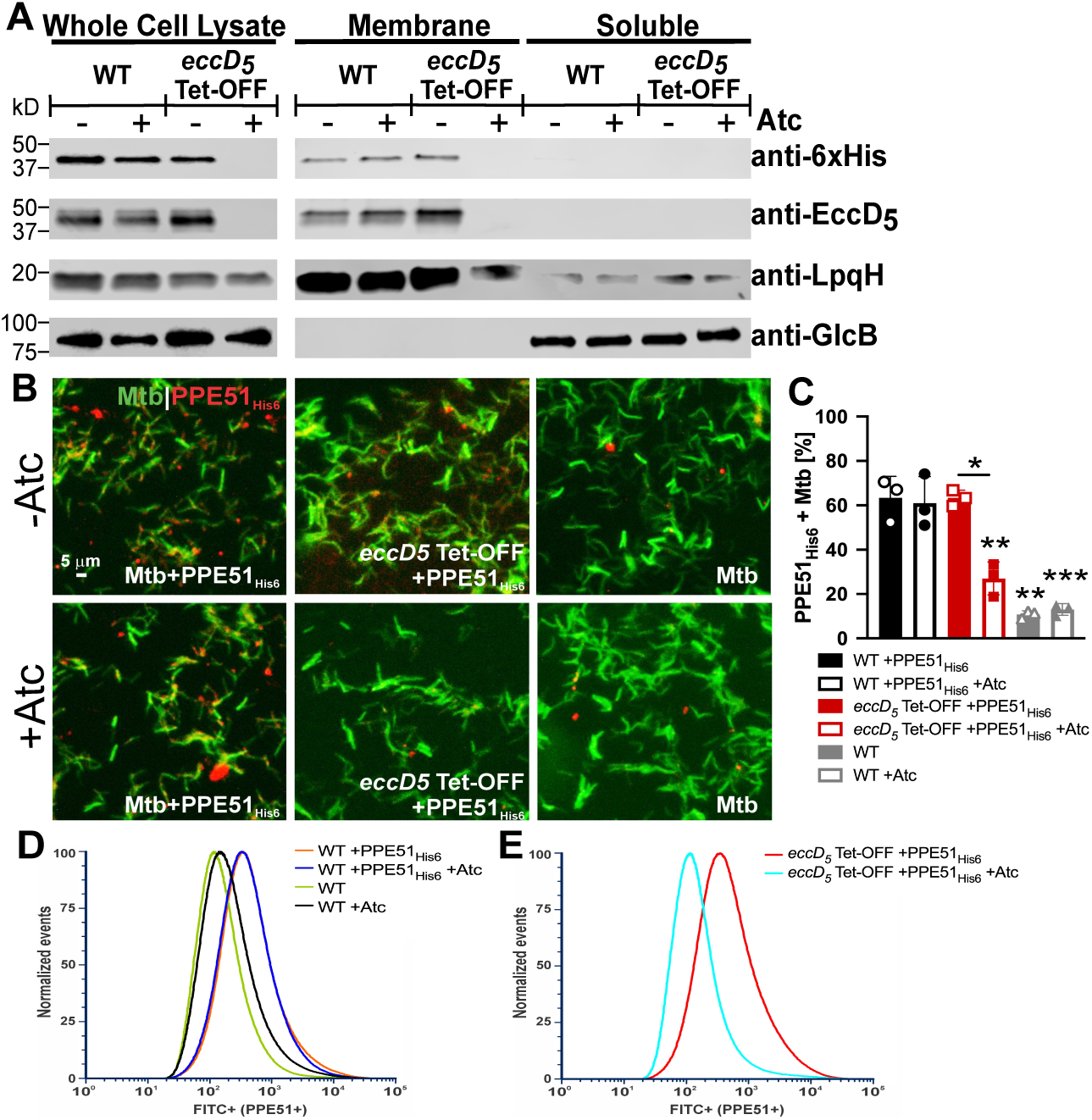
*M. tuberculosis* requires ESX-5 for outer membrane localization of PPE51. **(A**) Membrane localization of PPE51_His6_ by Western blotting. WT Erdman and *eccD_5_* Tet-OFF carrying pMV-*ppe51_His6_* were grown in Sauton’s medium ± 100 ng/ml anhydrotetracycline (Atc) to stationary phase for protein extraction. Whole cell lysates were separated into membrane and soluble fractions by ultracentrifugation. PPE51_His6_, EccD_5_, GlcB and LpqH were detected in equal amounts of total lysate, membrane and soluble fractions by Western blotting. GlcB and LpqH are soluble and membrane fraction loading controls, respectively. Results are representative of two independent experiments. (**B**) Detection of surface-accessible PPE51_His6_ in *M. tuberculosis* by fluorescence microscopy. The indicated *M. tuberculosis* strains were metabolically labeled with DMN-trehalose (green) and stained with monoclonal anti-6xHis antibody to detect PPE51_His6_ (red). (**C**) Percentage of cells scored positive for PPE51_His6_ relative to the untagged WT strain quantified from the images in **B** (n=3 technical replicates). Data are means ± standard deviations. Statistical analysis was performed between Atc (100 ng/ml) treated and untreated conditions for each strain (\**p*<0.05 \*\**p*<0.01; unpaired t-test). For each condition, statistical significance was calculated in comparison to WT *M. tuberculosis* with pMV- *ppe51_His6_* (\*\**p*<0.01, \*\*\**p*<0.001; one-way ANOVA with Dunnett’s correction). (**D-E**) Surface-accessible PPE51_His6_ determined by flow cytometry using an anti-6xHis antibody and Alexa fluor 488 conjugated secondary antibody in *M. tuberculosis* cells grown ±100 ng/ml Atc. (**D**) Overlay of WT pMV-*ppe51_His6_* and untagged WT ± Atc. (**E**) Overlay of *eccD_5_* Tet-OFF pMV-*ppe51_His6_* ± Atc.

To confirm that ESX-5 exports PPE51 to the *M. tuberculosis* outer membrane, we conducted antibody surface staining followed by both microscopy and flow cytometry. PPE51_His6_ was readily detected on the surface of WT Erdman and on *eccD_5_* Tet-OFF grown without Atc, which was reduced by EccD_5_ depletion with Atc (**Fig 3B**). Quantification revealed that Atc significantly reduced the percentage of *eccD_5_* Tet-OFF bacteria with detectable surface-localized PPE51_His6_ to the background level of WT *M. tuberculosis* lacking the PPE51_His6_ expression vector (**Fig 3C**). Analysis of 50,000 individual cells by flow cytometry also showed that PPE51_His6_ was readily detected on the surface of WT *M. tuberculosis*, as indicated by an increased fluorescence intensity relative to bacteria lacking the pMV-*ppe51_His6_* expression vector (**Fig 3D**). PPE51_His6_ was also detected on the surface of *eccD_5_* Tet-OFF grown without Atc, but not when EccD_5_ was depleted with Atc (**Fig 3E**). Collectively, these data indicate that *M. tuberculosis* exports PPE51 to the outer membrane via the ESX-5 system.

### *M. tuberculosis* requires ESX-5 to grow in and induce inflammatory cell death by macrophages

To determine the importance of ESX-5 during *M. tuberculosis* growth within phagocytes, we used the human THP-1 monocyte cell line differentiated to macrophages. To assess *M. tuberculosis* replication, THP-1 cells were infected at a low MOI (1:20 bacteria:macrophage) with WT or *eccD_5_* Tet-OFF that were pre-grown ± Atc for 3 days to deplete EccD_5_. Depletion of EccD_5_ significantly reduced *M. tuberculosis* growth in resting THP-1 cells, with over a 1 log decrease in CFU at day 7 (**Fig 4A**). *M. marinum* ESX-5 was implicated in activation of inflammatory cell death and IL-1β cytokine release in THP-1 cells (13). To test the if *M. tuberculosis* ESX-5 is similarly required to induce cell death, THP-1 cells were infected at a high MOI (1:1) and viable CFU, cytotoxicity and IL-1β release were assessed. EccD_5_ depletion +Atc did not significantly alter *M. tuberculosis* survival at 72 hr (**Fig 4B**). However, EccD_5_ depletion significantly reduced THP-1 cell death induced by *M. tuberculosis* infection (**Fig 4C**). In addition, THP-1 cells secreted significantly less IL-1β when infected with *M. tuberculosis* in which EccD_5_ was depleted (**Fig 4D**). Collectively, these data demonstrate that ESX-5 plays critical roles in replication of *M. tuberculosis* within resting phagocytes, which may be due its importance in nutrient uptake, and in stimulating inflammatory responses by infected cells.

**Figure 4.**
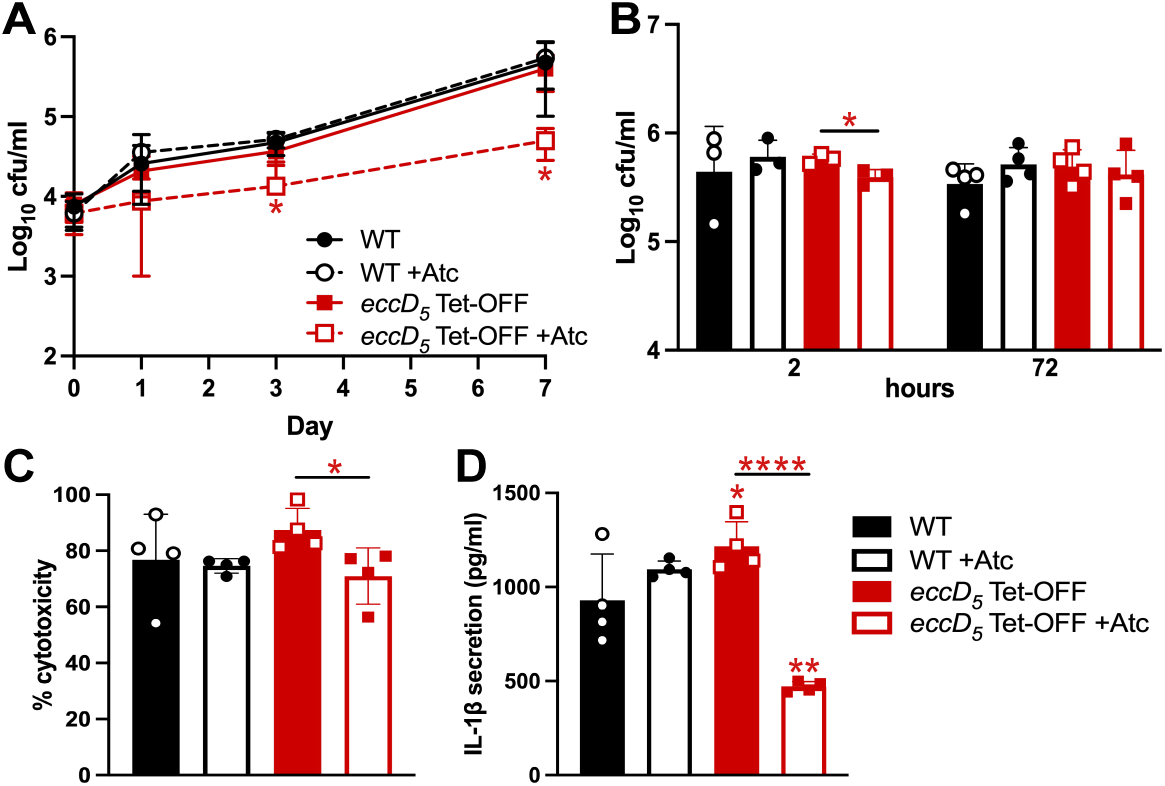
*M. tuberculosis* requires ESX-5 to grow in and induce inflammatory cell death of THP-1 cells. THP-1 cells were infected with the indicated *M. tuberculosis* strains grown in complete 7H9 ± 100 ng/ml anhydrotetracycline (Atc) and cultured ± 200 ng/ml Atc. (**A**) Cells were infected a low MOI (1:20, bacteria:macrophage). Viable *M. tuberculosis* were determined at the indicated times post-infection by plating. Data are means ± standard deviations of 3 independent experiments. Statistical analysis was performed between Atc treated and untreated conditions for each strain (\**p*<0.05, unpaired t-test). (**B-D**). THP-1 cells were infected at MOI 1:1 (bacteria:macrophage). (**B**) Viable *M. tuberculosis* were determined by plating at 2 and 72 hr. (**C**) Cytotoxicity was measured by a lactate dehydrogenase release assay. (**D**) Secreted IL-1β was measured by ELISA. Data are means ± standard deviations (n=4). (**B-D**) Statistical analysis was performed between Atc treated and untreated conditions (\**p*<0.05, \*\*\*\**p*<0.0001; unpaired t-test) and to compare each strain and condition to the WT no Atc control (\**p*<0.05, \*\**p*<0.001; one-way ANOVA with Dunnett’s correction).

### *M. tuberculosis* requires ESX-5 to replicate, disseminate and persist in aerosol-infected mice

To assess the role of ESX-5 in mammalian infection, we infected C57BL/6J mice by the aerosol route with either WT *M. tuberculosis* Erdman or *eccD_5_* Tet-OFF. We used a standard dose of doxycycline (dox, 2,000 ppm in chow) for conditional expression from Tet-regulated promoters in mice (49, 50), to deplete EccD_5_ at various times post-infection (**Fig 5A**). For WT Erdman, dox treatment modestly impaired replication and persistence in the lungs (**Fig 5B**) but did not significantly reduce dissemination to the spleen (**Fig 5C**). Without dox treatment, the *eccD_5_* Tet-OFF strain replicated in the lungs similarly to the WT control during acute infection, achieving a similar lung burden at 2 weeks post-infection (**Fig 5B**). The *eccD_5_* Tet-OFF strain also disseminated to the spleen similarly to the WT control (**Fig 5C**). However, during chronic infection (8-12 weeks post-infection), significantly fewer *eccD_5_* Tet-OFF bacteria were recovered from the lungs of mice as compared to the WT no dox control (**Fig 5B**). These data suggest that *eccD_5_* expression from the Tet-OFF promoter is sufficient for normal ESX-5 function during acute infection. However, unregulated expression of *eccD_5_* from the Tet-OFF promoter may either increase or decrease ESX-5 activity during chronic infection, leading to modest attenuation.

**Figure 5.**
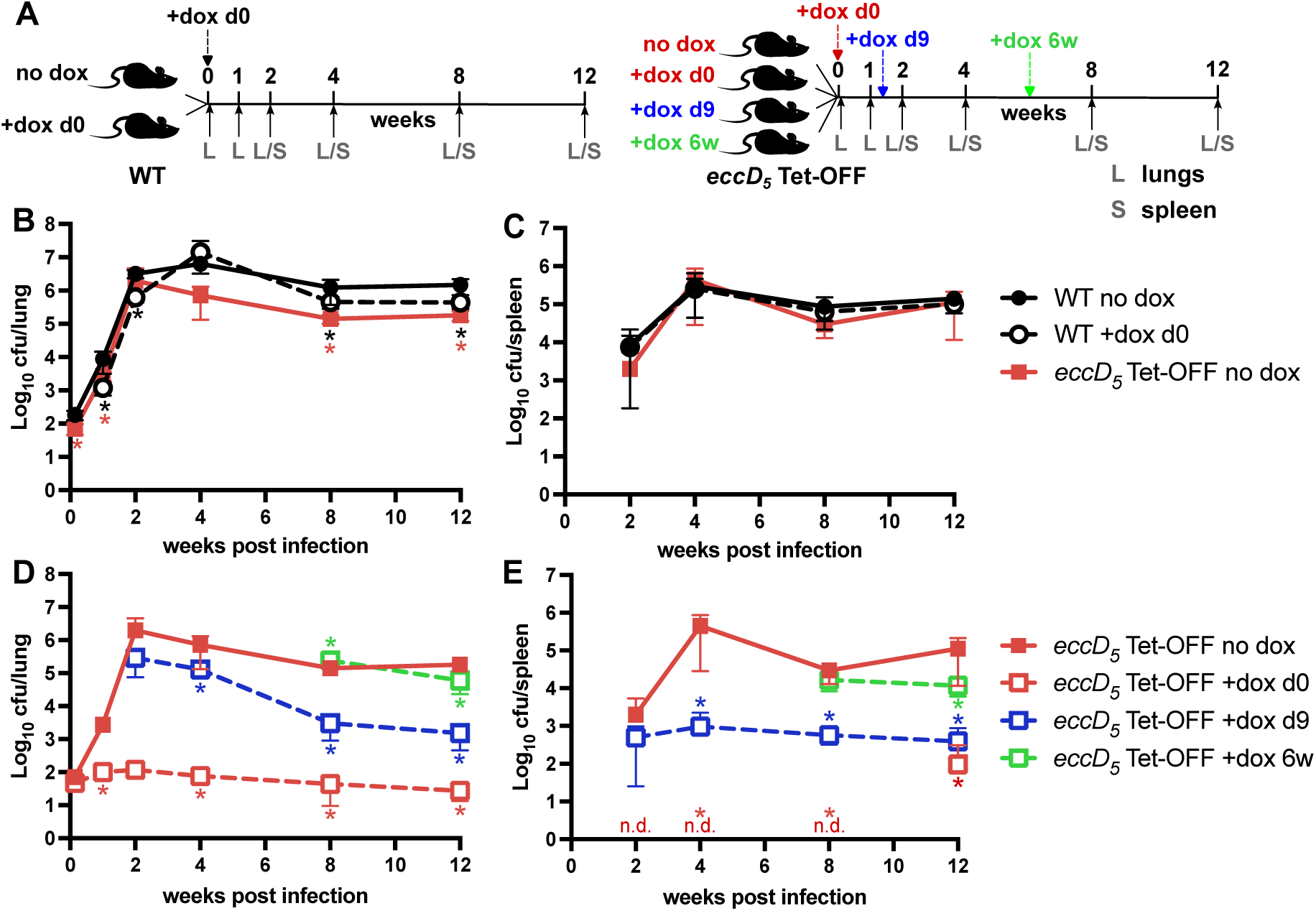
*M. tuberculosis* requires ESX-5 to replicate and disseminate in aerosol-infected mice. (**A**) Experiment design. Mice were infected with either WT Erdman or *eccD_5_* Tet-OFF by aerosol with ∼100 CFU. Mice were divided into groups that were either untreated or treated with doxycycline (dox) starting at day 0, day 9 or week 6. Groups of mice (n=6; 3 male, 3 female) were euthanized at the indicated times post-infection for collection of lung and spleen tissues. (**B-E**) Viable *M. tuberculosis* CFU determined by plating lung (**B, D**) or spleen (**C, E**) homogenates on 7H10 agar. Plates were incubated at 37°C for at least 4 weeks before counting CFU. Data are means ± standard deviations. Data for *eccD_5_* Tet-OFF no dox are reproduced in panels B, C and panels D, E to enable comparisons with both the WT controls and the + dox groups. Red n.d. in panel E indicates none detected for *eccD_5_* Tet-OFF +dox day 0 (limit of detection = 3 CFU). (**B, C**) Statistical analysis was performed by a one-way ANOVA with Tukey’s correction at each time point (\**p*<0.05). (**D, E**) A one-way ANOVA with Dunnett’s correction was used to compare each dox-treated group to the *eccD_5_* Tet-OFF no dox control (\**p*<0.05).

To determine at what stage(s) of infection *M. tuberculosis* requires ESX-5 activity, mice infected with the *eccD_5_* Tet-OFF strain were treated with dox starting at day 0 (initiation of infection), day 9 (acute infection) or week 6 (chronic infection) (**Fig 5A**). We compared the number of viable *eccD_5_* Tet-OFF bacteria in dox-treated mice to the no dox control. Treatment with dox starting at day 0 prevented replication of *eccD_5_* Tet-OFF bacteria in the lungs and dissemination to the spleens (**Fig 5D** and **5E**). The number of viable bacteria in the lungs increased slightly during the first two weeks of infection (from 47 CFU to 120 CFU, on average), but then slowly declined, reaching 28 CFU on average at 12 weeks post-infection (**Fig 5D**). While *eccD_5_* Tet-OFF bacteria were not detected in the spleens of dox-treated mice through 8 weeks post-infection, at 12 weeks post-infection we found viable CFU in the spleens of 3 of 6 mice (**Fig 5E**). We recovered only one colony from one of the mice, which may represent cross-contamination, but two other mice had similar CFUs in the spleens and lungs, suggesting late dissemination of *eccD_5_* Tet-OFF bacteria (**Fig 5E**). Both mice with high spleen CFU were male, but there was not any obvious trend of increased CFU of the *eccD_5_* Tet-OFF strain in male mice in other mouse groups (**Fig S4**).

To confirm that the *eccD_5_* Tet-OFF bacteria that disseminated to the spleen in dox-treated mice maintain Tet-repressible regulation of *eccD_5_*, we recovered colonies from the spleen of each mouse and tested *in vitro* growth and EccD_5_ protein production. Each *eccD_5_* Tet-OFF spleen isolate failed to grow on glycerol and glucose with Atc, similar to the parental control (**Fig S5A-F**). Each *eccD_5_* Tet-OFF spleen isolate also exhibited EccD_5_ depletion upon treatment with Atc (**Fig S5G**). In a Van10P vancomycin susceptibility assay for PDIM production (31), all isolates were equally resistant as the WT control (**Fig S5H**) suggesting they produce PDIM.

To determine if these strains harbored secondary mutations that enabled dissemination to the spleen, we conducted whole genome resequencing using both Illumina short-read and Oxford Nanopore long-read technologies on one isolate from each mouse. In the isolate from mouse 2, which had 525 CFU in the spleen, we did not identify any secondary mutations. We identified a 21 bp deletion in *ppe3* in the isolate from mouse 3 (42 CFU) and short deletions in two genes encoding PE_PGRS proteins in the isolate from mouse 6 [1 CFU; *Erdman_1299* (*rv1157c*), 7 bp; *Erdman_1942,* 10 bp] by short-read sequencing. However, long-read sequencing did not confirm mutations in these genes, suggesting they were errors due to the difficulty mapping short sequencing reads to the repetitive *pe* and *ppe* genes. Because we did not identify secondary mutations in *eccD_5_* Tet-OFF strains recovered from dox-treated mouse spleens, it is possible that dissemination of ESX-5 deficient bacteria occurs randomly due to a loss of lung containment.

To determine the importance of ESX-5 during acute infection, we initiated dox treatment of mice infected with *eccD_5_* Tet-OFF at day 9 post-infection (**Fig 5A**). We found that CFU of the *eccD_5_* Tet-OFF strain were significantly reduced at 4-12 weeks compared to the no dox control (**Fig 5D**). Even in the 5 days between initiation of dox treatment and the 2 week time point, we observed reduced replication of *eccD_5_* Tet-OFF bacteria in the lungs (**Fig 5D**) and reduced dissemination to the spleen (**Fig 5E**). After the 2 week time point, dox treatment caused clearance of *eccD_5_* Tet-OFF bacteria from the lungs (**Fig 5D**). However, we did not observe any reduction in *eccD_5_* Tet-OFF bacteria in the spleens over time with dox treatment initiated at day 9 (**Fig 5E**). At the dose we used, dox accumulates to a sufficient concentration in the spleen to fully repress expression from Tet-repressible promoters (50). These data suggest that *M. tuberculosis* requires ESX-5 activity for replication and survival within the lung during acute phase replication, but that ESX-5 may be dispensable for *M. tuberculosis* survival in the spleen.

To determine the role of ESX-5 during chronic infection, mice infected with *eccD_5_* Tet-OFF were treated with dox starting at 6 weeks post-infection (**Fig 5A**). We observed modest but statistically significant reductions in viable *eccD_5_* Tet-OFF bacteria recovered from the lungs and spleens of dox-treated mice at 12 weeks post-infection relative to the no dox controls (**Fig 5D** and **5E**). These data suggest that *M. tuberculosis* may have a reduced requirement for ESX-5 during chronic infection. Alternatively, *M. tuberculosis* may not need to produce new ESX-5 membrane complexes during chronic infection because it is in a slow growing state (51), resulting in relatively limited impact of *eccD_5_* transcriptional repression on chronic phase persistence.

## Discussion

*M. tuberculosis* requires ESX-5 for growth in standard culture conditions *in vitro*, which has complicated analysis of its roles during infection. Here using conditional expression of the EccD_5_ membrane component, we show that ESX-5 has broad functions in *M. tuberculosis* physiology and pathogenesis. We demonstrate that *M. tuberculosis* requires ESX-5 for growth on multiple carbon sources *in vitro*, including glycerol and glucose that are the carbon sources in standard Middlebrook media, which may explain its essentiality. We show that PPE51 is a substrate of ESX-5 in *M. tuberculosis*. Failure to export PPE51, which is required for glucose and glycerol uptake (28), likely explains why EccD_5_-depleted *M. tuberculosis* cannot grow on these carbon sources. We also show that expression of the outer membrane porin MspA enables growth of EccD_5_-depleted *M. tuberculosis* on glycerol, suggesting that ESX-5 functions to export nutrient transporters like PPE51 to the outer membrane. We demonstrate that EccD_5_ depletion prevents *M. tuberculosis* replication in resting macrophages and in the lungs of mice. Importantly, depletion of EccD_5_ causes *M. tuberculosis* clearance from mouse lung tissues, demonstrating that ESX-5 activity is critical for pathogenesis. Determining whether ESX-5 is important for nutrient acquisition or exporting/secreting “effector” proteins that directly manipulate host cell functions during infection will require identifying and characterizing additional ESX-5 substrates.

Our data corroborate a recent report that *M. marinum* exports PPE51 via ESX-5 (52). Our data conclusively demonstrate by surface accessibility experiments that ESX-5 exports PPE51 to the outer membrane in *M. tuberculosis*. We detected PPE51 exclusively in the membrane fraction, which is consistent with prior studies in *M. tuberculosis* (28). In *M. marinum*, PPE51 was also detected in the soluble fraction and secreted in culture filtrates, so it may not be efficiently retained in the *M. marinum* outer membrane (52). Our data in *M. tuberculosis* support a role for ESX-5 in activating macrophage inflammatory responses, including production of IL-1β, which was previously described in *M. marinum* (13). However, an ESX-5 deficient *M. marinum* strain was hyper-virulent in adult zebrafish (53), which contrasts with strongly reduced virulence of EccD_5_-deficient *M. tuberculosis* in mice. These differences highlight the importance of characterizing ESX-5 secretion system functions in *M. tuberculosis*, in addition to model organisms like *M. marinum*.

Our data also show that the *M. smegmatis* porin MspA cannot fully rescue growth of ESX-5-deficient *M. tuberculosis* in media with glycerol as the carbon source, which contrasts with *M. marinum* in which expression of MspA enabled deletion of either *eccC_5_* or *mycP_5_*. These data imply that *M. tuberculosis* ESX-5 has functions that are distinct from the function of ESX-5 in *M. marinum*. MspA fully rescued growth of EccD_5_-deficient *M. tuberculosis* in medium with trace heavy metals, suggesting that ESX-5 is required for metal resistance. Our data suggest that Cu^2+^ and/or Zn^2+^ could be toxic to ESX-5-deficient *M. tuberculosis*. Cu^2+^ and Zn^2+^ are concentrated in macrophage phagosomes and restrict *M. tuberculosis* replication (46, 47, 54). While ATPases that pump Cu^2+^ and Zn^2+^ across the *M. tuberculosis* inner membrane have been identified (46, 55, 56), mechanisms of metal efflux across the outer membrane are unknown. Metal efflux may be facilitated by PE and/or PPE protein substrates of the ESX-5 system.

While *M. tuberculosis* requires ESX-5 to export PPE51 and use certain carbon sources *in vitro*, the growth defects associated with EccD_5_ depletion in mice are unlikely to be due exclusively to reduced PPE51 export. *M. tuberculosis* does not require catabolism of glycerol or glucose to replicate in the lungs of acutely infected mice and mutants that cannot use glucose are only modestly attenuated during chronic infection (57, 58). ESX-5 may be required during acute infection to enable use of fatty acids that are considered primary carbon sources in infected cells (59). EccD_5_-depleted *M. tuberculosis* likely fails to grow on the model fatty acid ester Tween-40 due to reduced export of lipase or esterase enzymes that hydrolyze Tween-40 to release free fatty acids that are taken up by Mce1 (60). Many PE and PPE proteins have lipase or esterase activity (61-63) and are likely to be exported to the outer membrane by ESX-5. Several PE and PPE proteins have also been implicated in acquisition of heme iron (37, 38).

Our future studies will include identification of additional ESX-5 substrates that are required to use other nutrients, including carbon sources and heme, and determining the importance of ESX-5 dependent export of these substrates during infection. Our data show that *M. tuberculosis* requires ESX-5 for replication in the lungs and dissemination to the spleen in acutely infected mice, suggesting that ESX-5 is critical for host-pathogen interactions from the earliest stage of infection. The *M. tuberculosis* ESX-3 secretion system is similarly required during acute infection, as a Δ*esx-3* mutant was rapidly cleared from the lungs and failed to disseminate to the spleen in aerosol-infected C57BL/6 mice (9). *M. tuberculosis* ESX-3, like ESX-5, is essential for growth in standard culture conditions *in vitro* due to a role in iron acquisition (9). In contrast, *M. tuberculosis* mutants lacking ESX-1, which is considered a primary virulence factor, replicate in the lungs and spleens of C57BL/6 mice, but at a slower rate than WT bacteria (64-66). Our data suggest that ESX-5 is at least as important in *M. tuberculosis* pathogenesis as these other ESX secretion systems and support further characterization of its functions during infection.

We were surprised that EccD_5_ depletion at the initiation of infection did not cause clearance of *M. tuberculosis* from mouse lungs. We did not deplete EccD_5_ before infecting the mice, so the bacteria may remain viable due to pre-existing ESX-5 secretion complexes, which are not disrupted by dox treatment that only represses *eccD_5_* transcription. At this early stage of infection, *eccD_5_* transcriptional repression may restrict *M. tuberculosis* within resting alveolar macrophages. EccD_5_ depletion limited production of the inflammatory cytokine IL-1β by infected THP-1 cells. Infected alveolar macrophages must produce IL-1β to enable their migration from the airways to the lung interstitium (67). We speculate that because ESX-5 deficient *M. tuberculosis* fails to induce IL-1β, infected alveolar macrophages do not migrate into the lung tissue and therefore fail to activate an adaptive immune response. We observed dissemination to the spleen in a subset of animals at a late time point in these mice, which we could not attribute to any secondary mutations. Alveolar macrophages are very long-lived in mice, with a life-span of at least several months (68). When these infected cells die, *M. tuberculosis* may be taken up by other cell types that can escape from the lung to enable dissemination to the spleen.

Importantly, we observed clearance of EccD_5_-depleted *M. tuberculosis* from the lungs of mice that were treated with dox during acute infection. We observed clearance after 2 weeks post-infection, which coincides with the time when T cells that produce IFN-γ are recruited to the lungs (69). ESX-5 may be particularly important for *M. tuberculosis* to resist stressors in activated macrophages, such as metal toxicity. While dox treatment caused clearance of *eccD_5_* Tet-OFF bacteria from the lungs, it only limited initial dissemination to the spleen. We do not know the basis for persistence of *eccD_5_* Tet-OFF *M. tuberculosis* in the spleen during dox treatment, but it is unlikely to be related to the dox concentration, which is high enough to repress the Tet-OFF promoter (50). *M. tuberculosis* may infect a different cell type in the spleen, in which it does not require ESX-5 activity. Alternatively, *M. tuberculosis* may enter a slowly replicating state in the spleen, in which it can persist using pre-existing ESX-5 secretion complexes. This may also explain why *eccD_5_* transcriptional repression during chronic infection caused only modest reductions in bacterial burdens, as *M. tuberculosis* replicates slowly in this phase of infection (51). Alternatively, ESX-5 activity could be less important during chronic infection if it primarily enables *M. tuberculosis* growth by exporting outer membrane transporters or enzymes for nutrient acquisition. Distinguishing between these possibilities will require either using dual control systems for conditional protein degradation of ESX-5 components, which may be possible for the cytosolic EspG_5_ chaperone and EccA_5_ ATPase, or developing small molecule inhibitors of ESX-5.

Finally, our data indicate that ESX-5 is a strong candidate for development of new anti-tubercular drugs. Small molecule screens identified inhibitors of *M. tuberculosis* ESX-1 which act as anti-virulence drugs and limit bacterial growth in cell culture infection models (70-72). Inhibitors of ESX-5 might be more effective, as loss of ESX-5 function prevents *M. tuberculosis* growth and leads to clearance from infected mice. A screen for small molecules that inhibit *M. marinum* ESX-5 was reported, but the compounds identified caused dysregulated export of the LipY lipase reporter and did not specifically inhibit ESX-5 (73). Similar screens to those reported for ESX-1 could be done to identify small molecule inhibitors of *M. tuberculosis* ESX-5. Structures of the ESX-5 inner membrane complex exist to guide drug development (14, 15, 74), but the mechanisms by which ESX-5 substrates become incorporated into or pass through the *M. tuberculosis* outer membrane remain unknown. Identifying the ESX-5 outer membrane components will be important, as small molecules targeting these components would not need to pass through the outer membrane permeability barrier to have activity. Our future studies will seek to define these ESX-5 outer membrane components and determine how they mediate ESX-5 substrate export.

## Materials and Methods

### Bacterial strains and culture conditions

WT *M. tuberculosis* Erdman, *eccD_5_* Tet-OFF (19) and derivative strains (**Table S1**) were grown at 37°C with aeration in Middlebrook 7H9 medium (BD Difco) supplemented with 10% albumin-dextrose-saline (ADS), 0.5% glycerol and 0.1% Tween-80 (complete 7H9) or on Middlebrook 7H10 agar (BD Difco) supplemented with 10% oleic acid-albumin-dextrose-catalase (OADC; BD Biosciences) and 0.5% glycerol, unless otherwise noted. Frozen stocks were prepared by growing cultures to late-exponential phase and adding glycerol to 15% final concentration and were stored at -80°C. Antimicrobials were used at the following concentrations: kanamycin (Kan) 25 μg/ml for agar or 15 μg/ml for liquid, hygromycin (Hyg) 50 μg/ml, and cycloheximide 100 μg/ml.

### Growth curves

Bacteria were grown to mid-exponential phase (OD_600_ of ∼0.5) in complete 7H9 medium, then diluted to OD_600_ = 0.05 in fresh complete 7H9 medium ± 100 ng/ml anhydrotetracycline (Atc, Sigma) for three days to pre-deplete EccD_5_. Bacteria were collected by centrifugation (10 min, 2850 x*g*) washed twice in PBS with 0.01% tyloxapol (Sigma), then inoculated in triplicate at OD_600_ = 0.01 in Middlebrook 7H9 base (BD Difco) containing 0.01% tyloxapol and a sole carbon source: 0.5% glycerol, 0.2% glucose, 0.2% Tween-40 (Sigma), or 0.01% cholesterol (Sigma). Cholesterol stocks were prepared at 100 mg/ml in a 1:1 EtOH:tyloxapol solution at 80°C as described (75). Fresh cholesterol was added to 0.01% final concentration every 2-3 days from a cholesterol stock pre-warmed to 80°C. To test growth in minimal salts medium, bacteria grown in complete 7H9 medium ± 100 ng/ml Atc to pre-deplete EccD_5_ were washed twice in minimal medium without a carbon source (1 g/L KH_2_PO_4_, 2.5 g/L Na_2_HPO_4_, 0.5 g/L (NH_4_)_2_SO_4_, 0.15 g/L asparagine, 50 mg/L ferric ammonium citrate, 0.5 g/L MgSO_4_x7H_2_O, 0.5 mg/L CaCl_2_, 0.1 mg/L ZnSO_4_, 0.05% tyloxapol), then diluted in duplicate at OD_600_ = 0.05 in minimal medium with no added carbon, 0.2% glycerol or 0.2% glucose ± 100 ng/ml Atc. To test heavy metal toxicity, bacteria were grown in homemade 7H9 (2.5 g/L Na_2_HPO_4_, 1.0 g/L KH_2_PO_4_, 0.5 g/L monosodium glutamate, 0.5 g/L (NH_4_)_2_SO_4_, 0.05 g/L MgSO_4_x7H_2_O, 1.0 mg/L pyridoxine HCl, 0.5 mg/L biotin, 0.5 mg/L CaCl_2_) without added Fe^3+^, Cu^2+^ or Zn^2+^. Fresh Atc (100 ng/ml) was added to cultures every 7 days to maintain *eccD_5_* transcriptional repression. The OD_600_ of cultures was measured every 2-3 days. For minimal medium, cultures were serially diluted and plated on 7H10 agar at days 0, 7 and 14. Plates were incubated at 37°C for 3-4 weeks before counting CFU.

### Cloning and strain construction

Plasmids used in this study are listed in **Table S2**. The pMV- *mspA* vector was generated by cloning *M. smegmatis mspA* and the mycobacterial optimal promoter *P_smyc_* from pMN016 (27) into the episomal vector pMV261. The pMV261 vector was digested with XbaI and ClaI to remove the P*_hsp60_* promoter. *M. smegmatis mspA* and *P_smyc_* were removed from pMN016 by digestion with XbaI and ClaI. The digested products were gel purified (QIAquick gel extraction; Qiagen), ligated with T4 DNA ligase (NEB) and transformed in *E. coli* DH5α. The pMV-*mspA* vector was confirmed by XbaI/ClaI digest and Sanger sequencing.

The pMV-*ppe51_His6_* vector was made by cloning *ppe51_His6_* from the integrating vector pMV306-NHK-*ppe51_His6_* (28) into pMV261. Plasmids were digested with XbaI and HindIII, the pMV261 vector and *ppe51_His6_* insert were gel purified (QIAquick gel extraction; Qiagen), ligated with T4 DNA ligase and transformed in *E. coli* DH5α. The pMV-*ppe51_His6_* vector was confirmed by XbaI/HindIII digest and Sanger sequencing.

The pMV261, pMV-*mspA*, and pMV-*ppe51_His6_* plasmids were transformed into WT Erdman and *eccD_5_* Tet-OFF by electroporation as described (76). Transformants were selected by plating on 7H10 with Kan (WT) or 7H10 with Kan and Hyg (*eccD_5_* Tet-OFF) and confirmed by PCR using primers listed in **Table S3**.

### Protein extraction, fractionation and Western blotting

*M. tuberculosis* strains were grown in complete 7H9 medium to mid-exponential phase, washed twice in Sauton’s medium (4 g/L L-Asparagine, 0.5 g/L potassium phosphate monobasic, 0.5 g/L MgSO_4_x7H_2_O, 2 g/L citric acid, 0.05 g/L ferric ammonium citrate, 60 ml/L glycerol, 1 mg/L ZnSO_4_) with 0.1 % Tween-80 and resuspended at OD_600_ = 0.2 in 30 ml Sauton’s +0.1% Tw-80 ± 100 ng/ml Atc. Cultures were grown at 37°C to late-logarithmic phase (OD_600_ ∼1.2). Bacteria were pelleted (2850 x*g*, 10 min), resuspended in 30 ml Sauton’s medium without detergent, then grown with aeration for 6 days at 37°C. Bacteria were collected by centrifugation (4100 xg, 15 min, 4°C) and washed twice in ice-cold PBS. Cell pellets were resuspended in 1 ml ice-cold PBS containing Complete EDTA-free protease inhibitors (Roche) and bacteria were lysed by bead beating. Large cell debris was removed by centrifugation and lysates were filter sterilized with a 0.2 μm Nanosep MF centrifugal filter (Pall Life Sciences) as described (77). Whole cell lysates were separated into membrane and soluble fractions by two rounds of ultracentrifugation (100,000 x*g*, 1 hr, 4°C) as described (37). The final membrane pellet was resuspended in PBS with protease inhibitors and 1% SDS.

Protein concentrations in whole cell lysates and soluble fractions were determined by BCA assay (Pierce). Equal amounts of total protein or equivalent volumes of the membrane fraction based on whole cell lysate protein concentrations were separated by SDS-PAGE on AnyKD gels (BioRad) and transferred to 0.2 μm nitrocellulose membranes. Membranes were blocked in PBS with 0.1% Tween-20 (PBST-20) with 5% nonfat milk powder for 1 hr, washed with PBST-20 and probed overnight at 4°C with primary antisera diluted in PBST-20 with 2.5% nonfat milk powder. Primary antisera were used at the following dilutions rabbit anti-6xHis 1:5,000 (ZooMab ZRB2297, Sigma-Aldrich), rabbit anti-EccD_5_ 1:1,000 (19), mouse anti-LpqH 1:1,000 (NR-13792, clone IT-54, BEI Resources), mouse anti-GlcB 1:500 (NR-13799, BEI Resources), or mouse anti-GroEL2 1:1,000 (NR-13657, clone IT-70, BEI Resources). Membranes were washed with PBST-20, then probed for 1 hr with horseradish peroxidase conjugated secondary antibody (goat anti-rabbit IgG 31466 or goat anti-mouse IgG 31431, ThermoFisher) at 1:10,000 dilution in PBST-20 with 2.5% nonfat milk powder. Membranes were washed with PBST-20, incubated for 5 min with SuperSignal Pico West chemiluminescent substrate (ThermoFisher) and immediately imaged using an Odyssey Fc Imaging System (LI-COR) with LI-COR Image Studio software.

### Detection of surface-accessible PPE51_His6_ in *M. tuberculosis* by flow cytometry

Surface detection of PPE51_His6_ in *M. tuberculosis* Erdman and derivative strains was performed as previously described (3). Bacteria were grown in Middlebrook 7H9 supplemented with 10% OADC, 0.5% glycerol and 0.02% tyloxapol. To maintain PDIM production, 0.1 mM sodium propionate was added to the medium (31). Once the cultures reached an OD_600_ of ∼0.8, bacteria were pelleted and washed once in Sauton’s medium containing 0.02% tyloxapol. The cells were then cultured in the same medium with the required antibiotics until an OD_600_ of ∼1.0. Cultures were passed through a 5 μm syringe filter (Millipore) and allowed to grow until an OD_600_ of ∼1.0. Cells were fixed with 4% paraformaldehyde for 1 hr and blocked in 2.5% normal goat serum for 30 min. Subsequently, they were incubated with anti-His Tag antibody (1:50, ABclonal #AE003) for 1 hr followed by incubation with AlexaFluor 488-conjugated goat anti-mouse IgG (H+L) secondary antibody (1:500, Invitrogen) for another hour. Between each step, bacteria were washed three times with 1x PBS. For each sample, an unstained control (autofluorescence) was included, and 50,000 events were acquired per sample using a flow cytometer (LSRFortessa, BD Biosciences). Data were exported as FCS files, analyzed with FCS Express 7 software, and surface accessible PPE51_His6_ was displayed as histograms.

### Analysis of surface accessible PPE51_His6_ in *M. tuberculosis* by fluorescence microscopy

*M. tuberculosis* strains were cultured as for flow cytometry experiments. After passage through the 5 μm syringe filter (Millipore), bacteria were grown in Sauton’s medium containing 0.02% tyloxapol until OD_600_ of ∼0.8. Cells were then stained with 100 μg/ml DMN trehalose (78) and incubated overnight at 37°C. Following staining, cells were fixed with 4% paraformaldehyde for 1 hr, blocked with 2.5% normal goat serum for 30 min, and sequentially incubated with anti-His Tag antibody (1:50, ABclonal #AE003) and Alexa Fluor 594-conjugated goat anti-mouse IgG (1:300, Invitrogen), each for 1 hr. Bacteria were washed three times with PBS between each step. After staining, bacteria were smeared on coverslips and mounted on glass slides using ProLong Glass Antifade Mounting (Invitrogen). The samples were then analyzed using fluorescence microscope / Cytation 5. Image analysis was done with Gen5 software. A primary mask was applied based on GFP fluorescence (representing *M. tuberculosis*) with object size thresholds set between 2-10 μm. PPE51_His6_ expression was visualized using a TRITC filter and this fluorescence was used as the secondary mask. Mean TRITC intensity was measured within the primary mask. Scatter plots were generated in Gen5 and subpopulations with elevated fluorescence were gated relative to untagged wild-type *M. tuberculosis*. Bar graphs were plotted from the data obtained from Cytation 5 analysis using GraphPad Prism.

### Growth in THP-1 macrophages

The THP-1 (ATCC TIB-202) human monocytic cell line was cultured in RPMI base (ATCC or Gibco) supplemented with non-heat treated 10% fetal bovine serum (Corning or Gibco) (hereafter referred to as RPMI) in a humidified incubator at 37°C with 5% CO_2_. THP-1 cells were differentiated into monocyte-derived macrophages with 50 nM phorbol 12-myristate-13-acetate (PMA, Sigma) for 24 hr in 96-well tissue culture plates for experiments at 1:20 MOI or 24-well tissue culture plates for experiments at 1:1 MOI. Differentiated cells were washed twice with Hanks Balanced Salt Solution (Corning), given fresh RPMI, and rested for 72 hr before infection.

Bacteria were grown from frozen stocks in complete 7H9 to mid-exponential phase (OD_600_ 0.4-0.7), then diluted to OD_600_ = 0.05 in complete 7H9 ± 100 ng/ml Atc and grown for 72 hr prior to the infection. Bacteria were pelleted by centrifugation (2,850 x*g*, 10 min), resuspended in PBS containing 0.05% Tween-80 (PBS-T), declumped by centrifugation (58 x*g*, 5 min) and diluted in RPMI at the appropriate density for infection. The THP-1 cell culture media was refreshed with RPMI ± 200 ng/ml Atc. Infections were initiated at MOI = 1:20 (bacteria:macrophage) or MOI = 1 for 2 hours at 37°C with 5% CO_2_. Extracellular bacteria were removed by three washes with pre-warmed Dulbecco’s PBS (Gibco), RPMI ± 200 ng/ml Atc was replaced, and cells were incubated at 37°C with 5% CO_2_. For experiments at MOI = 1:20, viable intracellular bacteria were enumerated at 2 hr, 1, 3 and 7 days post-infection. For experiments at MOI = 1, bacteria were enumerated at 2 and 72 hr post infection. THP-1 cells were lysed at 37°C for 10 min with 0.1% Tween-80. Monolayers were visually inspected for lysis before serially diluting the lysate in PBS-T and plating on 7H10 to recover viable bacteria. Plates were incubated for at least 3 weeks at 37°C before counting CFU.

### Cell death and cytokine release assays

PMA-differentiated THP-1 cells infected at an MOI of 1 were assessed for cellular necrosis by measuring lactate dehydrogenase (LDH) release and for production of the inflammatory cytokine IL-1β at 3 days post-infection. Cell culture supernatants from infected cells and uninfected controls were filtered sterilized with a 0.2 μm Nanosep MF centrifugal filter (Pall Life Sciences; 14,000 x*g*, 3 min, 4°C). Supernatants were aliquoted and frozen at -80°C until use. LDH was measured with a CytoTox 96® Non-Radioactive Cytotoxicity Assay kit (Promega). Uninfected THP-1 control cells were lysed with 1x lysis solution provided with the kit for 45 min at 37°C to determine the maximal LDH release. Percent cytotoxicity was calculated as [LDH in culture supernatants (OD_492_)/maximal LDH from lysed uninfected cells (OD492)] x100. Human IL-1β was quantified in culture supernatants by ELISA (Invitrogen).

### Phthiocerol dimycocerosate (PDIM) assays

To directly measure PDIM production, *M. tuberculosis* cultures grown in 10 ml complete 7H9 medium to mid-logarithmic phase were labeled for 48 hours with 10 μCi of [1-^14^C] propionic acid, sodium salt (American Radiolabeled Chemicals, Inc; specific activity 50-60 mCi/mmol) prior to extraction of apolar lipids as described (25). Labeled lipids were separated by thin layer chromatography and detected by phosphor imaging as described (79). A Van10P assay was also used to indirectly assess PDIM production based on vancomycin (Van) susceptibility in the presence of propionate (31). Bacteria grown to late-exponential phase (OD_600_ ∼0.8) in complete Middlebrook 7H9 were diluted 1:100 in 7H9 containing 0.1 mM propionate ± 10 μg/ml Van. The OD_600_ was measured at 14 days and Van10P % growth was calculated as (Van10-P OD_600_/Van0-P OD_600_) x100.

### Mouse infections

Male and female C57BL/6J mice 6 weeks of age were purchased from the Jackson Laboratory, USA and infected by the aerosol route with ∼100 CFU of *M. tuberculosis* Erdman or *eccD_5_* Tet-OFF using an Inhalation Exposure System (GlasCol) as described (80). Bacteria grown to mid-exponential phase (OD_600_ of ∼0.5) were washed once with phosphate buffered saline with 0.05% Tween-80 (PBS-T) and diluted to OD_600_ = 0.007 in PBS-T for aerosol infection. Groups of mice were treated with doxycycline (dox) chow (irradiated Teklad Global 2018 rodent diet containing 2,000 ppm dox; Envigo) starting at day 0 (immediately after infection), day 9, or day 42 (week 6). Dox chow was stored at 4°C until use and was replaced weekly. Groups of mice (*n*=6; 3 male, 3 female) were euthanized by CO_2_ overdose at 24 hr and 1, 2, 4, 8 and 12 weeks post-infection to recover lung and spleen tissues. Organs were homogenized in PBS-T and bacterial CFU were determined by plating serially diluted homogenates on 7H10 containing cycloheximide. Colonies were counted after at least 4 weeks of incubation at 37°C.

### Ethics statement

All animal protocols used in this study were reviewed and approved by the University of Minnesota Institutional Animal Care and Use Committee (IACUC) under protocol number 2102-38860A. The University of Minnesota’s NIH Animal Welfare Assurance number is D16-00288 (A3456-01), most recent renewal date: 5/1/2024, expiration date: 4/30/2028. All animal experiments followed the recommendations established in the Guide for the Care and Use of Laboratory Animals of the National Institutes of Health (81).

### Whole genome sequencing

Genomic DNA was extracted from *M. tuberculosis eccD_5_* Tet-OFF and single *eccD_5_* Tet-OFF isolates from the spleens of mice treated with dox starting at day 0. Strains were grown to late-logarithmic phase (OD_600_ = 1.0) and DNA was extracted by the CTAB-lysozyme method (82). DNA was cleaned with the Genomic DNA clean and concentrator kit 25 (Zymo) and submitted to Microbial Genome Sequencing Center (MiGS, now SeqCenter, Pittsburgh, PA; *eccD_5_* Tet-OFF) or SeqCoast Genomics (Portsmouth, NH; *eccD_5_* Tet-OFF spleen isolates) for library preparation and Illumina sequencing. MiGS conducted library preparation with the Illumina DNA Prep kit and IDT 10 bp UDI indices and sequencing on an Illumina NextSeq 2000 (2x151 bp reads). Demultiplexing, quality control and adaptor trimming were conducted using bcl-convert (v3.9.3). SeqCoast conducted library preparation with the Illumina DNA Prep tagmentation kit and unique dual indices and sequencing on the Illumina NextSeq 2000 using a 300 cycle flow cell kit (2x150 bp reads) with 1-2% PhiX control spiked in for optimal base calling. Read demultiplexing, trimming and run analytics were performed using DRAGEN v3.10.12. To generate a consensus sequence for each strain, paired reads were mapped to the *M. tuberculosis* Erdman reference genome (NC_020559.1) using the “map to reference” function in Geneious Prime 2021 software (Biomatters, Ltd.) as previously described (79). To identify single nucleotide polymorphisms (SNPs) or small insertions and deletions (indels), whole genome alignments were performed with the Erdman reference genome, the published Tischler WT Erdman genome (79) and the *eccD_5_* Tet-OFF strains using the “Align Whole Genomes” function with the default Mauve Genome parameters as described (79).

For Oxford Nanopore Technology (ONT) long-read sequencing, genomic DNA was extracted, cleaned and concentrated as described above. The DNA concentration was determined using PicoGreen DNA quantification at the University of Minnesota Genomics Center (UMGC) and 200 ng of total DNA was used for library preparation. DNA samples were barcoded with the Rapid Barcoding Kit v14 (ONT). Barcoded samples were pooled and cleaned using AMPure beads (Beckman Coulter). The pooled library was quantified with a Qubit fluorometer (Invitrogen) and the library size was determined using the Genomic DNA ScreenTape® (Agilent). Rapid adaptors were added to the library immediately before loading on the MinION flow cell (ONT) and run using a GridION instrument (ONT). Portions of the prepared library were run on the MinION flow cell three times, with flow cell washing between runs, until all pores were depleted. ONT sequencing reads were collected with MinKNOW software and converted into fastq files. ONT fastq files were mapped to the *M. tuberculosis* Erdman reference genome (NC_020559.1) using Minimap2.24 in Geneious Prime software with the default settings for ONT long-read technology. The contig generated for each *eccD_5_* Tet-OFF spleen isolate was used to confirm or refute potential mutations identified by analysis of Illumina short-read sequencing data.

## Data availability

All raw sequencing data from whole genome sequencing underlying the results reported are available in FASTA format at the NCBI Sequence Read Archive (BioProject PRJNA1348213).

## Statistical analyses

A student’s unpaired *t*-test (two-tailed) was used for pairwise comparisons between strains or between conditions *in vitro* (± Atc). One-way ANOVA with Tukey’s or Dunnett’s correction applied post-hoc were used for comparisons of multiple strains. Two-way ANOVA with Tukey’s correction applied post-hoc were used for comparison of multiple strains and conditions. *P* values were calculated in R (CRAN r-project.org) or in GraphPad Prism (GraphPad Software, Inc). *p* values < 0.05 were considered statistically significant.

## Acknowledgements

Portions of this work were completed with resources, including instrumentation and staff, at the University of Minnesota Imaging Centers (UIC; RRID: SCR_020997) and at the University of Minnesota Genomics Center (UMGC; RRID: SCR_012413). We thank Dr. Clifton Barry for providing the pMV306-NHK-*ppe51_His6_* plasmid and Taylor Crooks for assistance with statistical analyses. The following reagents were obtained through BEI Resources, NIAID, NIH: Monoclonal Anti-*Mycobacterium tuberculosis* LpqH (Gene Rv3763), IT-54 (produced *in vitro*), NR-13792; Monoclonal Anti-*Mycobacterium tuberculosis* GlcB (Gene Rv1837c), Clone α-GlcB (produced *in vitro*), NR-13799; Monoclonal Anti-*Mycobacterium tuberculosis* GroEL2 (Gene Rv0440), Clone IT-70 (DCA4) (produced *in vitro*), NR-13657. This work was supported by NIH grants R21AI125475 (ADT), R21AI163580 (ADT) and R01AI175106 (MN) and by a University of Minnesota Research and Innovation Office Grant-In-Aid Bridge Award (ADT).

## SUPPLEMENTARY MATERIALS

**Fig S1.** *M. tuberculosis eccD_5_* Tet-OFF strains produce phthiocerol dimycocerosate (PDIM). The indicated strains were grown in complete 7H9 medium and labeled for 48 hr with 10 μCi ^14^C propionate prior to extraction of lipids and analysis of PDIM production by thin-layer chromatography. The DIM A and DIM B forms of PDIM are indicated. (**A**) WT Erdman and *eccD_5_* Tet-OFF. (**B**) WT Erdman, *eccD_5_* Tet-OFF pMV261 and *eccD_5_* Tet-OFF pMV-*mspA*.

**Fig S2.** *M. tuberculosis* requires ESX-5 for *in vitro* growth in minimal medium with glycerol or glucose. WT Erdman and *eccD_5_* Tet-OFF were grown in complete Middlebrook 7H9 ± 100 ng/ml Atc to mid-exponential phase, then washed and diluted to OD_600_ = 0.05 in minimal salts medium with 0.05% tyloxapol and no added carbon (**A, D**), 0.2% glycerol (**B, E**), or 0.2% glucose (**C, F**). Fresh Atc (100 ng/ml) was added to +Atc cultures every 7 days. Growth was monitored by OD_600_ measurements every 2 days (**A-C**) or by plating serially diluted cultures on Middlebrook 7H10 agar every 7 days (**D-F**). Data are means ± standard deviations of 4 biological replicates from 2 independent experiments. Statistical analysis was performed to compare *eccD_5_* Tet-OFF ± Atc to WT by one-way ANOVA with Dunnett’s correction (\**p*<0.05, \*\**p*<0.01, \*\*\*\**p*<0.0001).

**Fig S3.** Western blotting and 7H9 with no added carbon growth curve controls for *M. tuberculosis* strains expressing MspA. **(A,B)** Western blots to confirm EccD_5_ depletion. Whole cell lysates were prepared from cultures grown in 7H9 + 0.5% glycerol (**A**) or in 7H9 with trace Fe^3+^, Cu^2+^ and Zn^2+^ + 0.5% glycerol (**B**) that are shown in Fig 2. EccD_5_ and GroEL2 were detected in 5.8 μg total protein (**A**) or 10 μg total protein (**B**) by Western blotting. (**C,D**) The indicated strains were grown in complete Middlebrook 7H9 ± 100 ng/ml Atc to mid-exponential phase, then washed and diluted in triplicate to OD_600_ = 0.01 in home-made Middlebrook 7H9 with trace Fe^3+^ and Cu^2+^ with 0.01% tyloxapol and no added carbon ± 100 ng/ml anhydrotetracycline (Atc). Growth was monitored by OD_600_ measurements every 2-3 days. Fresh Atc (100 ng/ml) was added to +Atc cultures every 7 days. Data are means ± standard deviations. Statistical analysis was done to compare each strain and condition to the WT pMV261 untreated control (**C**) or to the *eccD_5_* Tet-OFF pMV261 untreated control (**D**) (\**p*<0.05; two-way ANOVA with a simple effects model and Dunnett’s correction).

**Fig S4.** *M. tuberculosis* lung and spleen burdens from individual mice, separated by sex. Data from C57BL/6 mice that were infected by aerosol with WT Erdman (**A, B, D, E**) or *eccD_5_* Tet-OFF (**C, F, G-L**) and treated with doxycycline (2,000 ppm in chow) starting at the indicated time points, from Fig 5. *M. tuberculosis* CFU in lung (**A-C, G-I**) and spleen (**D-F, J-L**) tissues were determined by plating and data from male and female mice (*n*=3 each) are shown separately. Lines indicate the mean. In **J**, n.d. indicates none detected in either male or female mice (detection limit = 3 CFU). Asterisks indicate statistically significant differences between male and female mice in each experimental group (\**p*<0.05; unpaired t-test).

**Fig S5.** *M. tuberculosis eccD_5_* Tet-OFF isolates recovered from spleens of dox-treated mice at 12 weeks post-infection retain Atc-repressible EccD_5_ expression. **(A-F)** *M. tuberculosis eccD_5_* Tet-OFF mouse spleen isolates or WT control were grown in complete Middlebrook 7H9 ± 100 ng/ml Atc to mid-exponential phase, then washed and diluted to OD_600_ = 0.01 in complete 7H9 (**A, C, E**), or modified 7H9 with 0.01% tyloxapol +0.5% glycerol +0.2% glucose (**B, D, F**) ± 100 ng/ml anhydrotetracycline (Atc). Growth was monitored by OD_600_ measurements every 3-4 days. Fresh Atc (100 ng/ml) was added to +Atc cultures at day 7. (**G**) *M. tuberculosis eccD_5_* Tet-OFF mouse spleen isolates and the WT control were grown in complete 7H9 ± 100 ng/ml Atc and whole cell lysates were prepared. EccD_5_ and LpqH were detected in 11.9 μg of total protein by Western blotting. (**H**) Vancomycin susceptibility in the presence of propionate quantified by Van10-P assay. Van10-P % growth is [(OD_600_ 10 μg Van)/(OD_600_ 0 μg Van)] x100.

**Table S1.** *M. tuberculosis* strains used in this study.

**Table S2.** Plasmids used in this study.

**Table S3.** Oligonucleotide primers used in this study.

