## Supplemental Tables and Figures for "The *Mycobacterium tuberculosis* ESX-5 secretion system enables carbon source utilization and growth in mice"

**Table S1. *Mycobacterium tuberculosis* strains used in this study.**

| **Strain name** | **Genotype** | **Source** |
|---|---|---|
| WT *M. tuberculosis* Erdman | wild type | Lab strain |
| *eccD5* Tet-OFF | Δ*eccD5* pGMCH-*eccD5* Tet-OFF, $Hyg^R$ | (1) |
| WT pMV261 | pMV261, $Kan^R$ | This study |
| WT pMV-*mspA* | pMV261-*mspA*, $Kan^R$ | This study |
| *eccD5* Tet-OFF pMV261 | Δ*eccD5* pGMCH-*eccD5* Tet-OFF pMV261, $Hyg^R$ $Kan^R$ | This study |
| *eccD5* Tet-OFF pMV-*mspA* | Δ*eccD5* pGMCH-*eccD5* Tet-OFF pMV-*mspA*, $Hyg^R$ $Kan^R$ | This study |
| WT pMV-*ppe51-His6* | pMV-*ppe51-His6*, $Kan^R$ | This study |
| *eccD5* Tet-OFF pMV-*ppe51-His6* | Δ*eccD5* pGMCH-*eccD5* Tet-OFF pMV-*ppe51-His6*, $Hyg^R$ $Kan^R$ | This study |

**Table S2. Plasmids used in this study.**

| **Plasmid name** | **Description** | **Source** |
|---|---|---|
| pMV261 | episomal plasmid with constitutive $P_{hsp60}$ promoter, $Kan^R$ | (2) |
| | episomal plasmid with *M. smegmatis mspA* under the $P_{smyc}$ optimal mycobacterial promoter | (3) |
| pMV-*mspA* | episomal plasmid with *M. smegmatis mspA* under the $P_{smyc}$ optimal mycobacterial promoter, $Kan^R$ | This study |
| pMV306-HNK-*ppe51-His6* | L5 *attB* integrating plasmid with *ppe51-His6* under the constitutive $P_{hsp60}$ promoter, $Kan^R$ | (4) |
| pMV-*ppe51-His6* | episomal plasmid with *ppe51-His6* under the constitutive $P_{hsp60}$ promoter, $Kan^R$ | This study |

**Table S3. Oligonucleotide primers used in this study.**

| **Primer name** | **Sequence 5'-3'** | **Use** |
|---|---|---|
| mspA_F | gagcacaggcacctctcac | Check for pMV-*mspA* in *M. tuberculosis* |
| mspA_R | gacgtcgaccgagaacgttg | Check for pMV-*mspA* in *M. tuberculosis* |
| pMV361_seqF | cagcgaggacaacttgagcc | Check for pMV-*ppe51-His6* in *M. tuberculosis* |
| ppe51R | ctcgcgagcaccgtgttg | Check for pMV-*ppe51-His6* in *M. tuberculosis* |

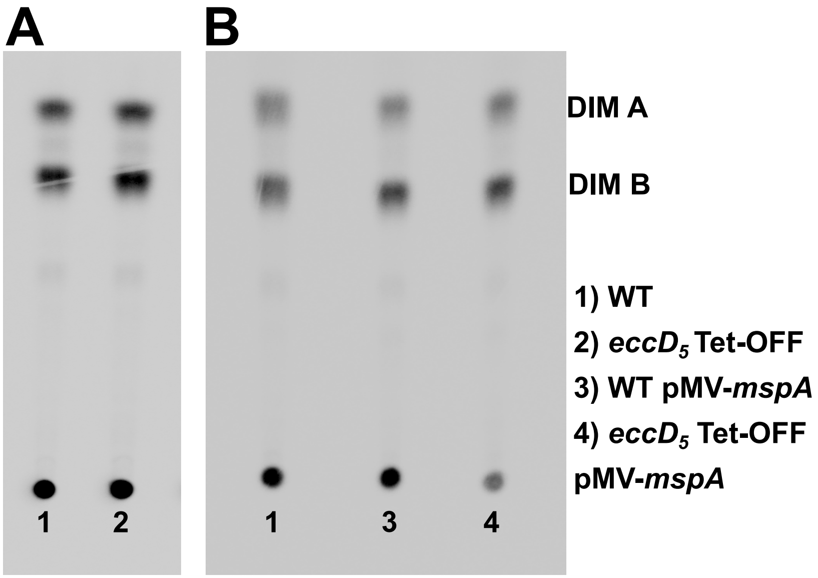


**Fig S1. *M. tuberculosis eccD*$_5$ Tet-OFF strains produce phthiocerol dimycocerosate (PDIM).** The indicated strains were grown in complete 7H9 medium and labeled for 48 hr with 10 μCi $^{14}C$ propionate prior to extraction of lipids and analysis of PDIM production by thin-layer chromatography. The DIM A and DIM B forms of PDIM are indicated. (**A**) WT Erdman and *eccD*$_5$ Tet-OFF. (**B**) WT Erdman, *eccD*$_5$ Tet-OFF pMV261 and *eccD*$_5$ Tet-OFF pMV-*mspA*.

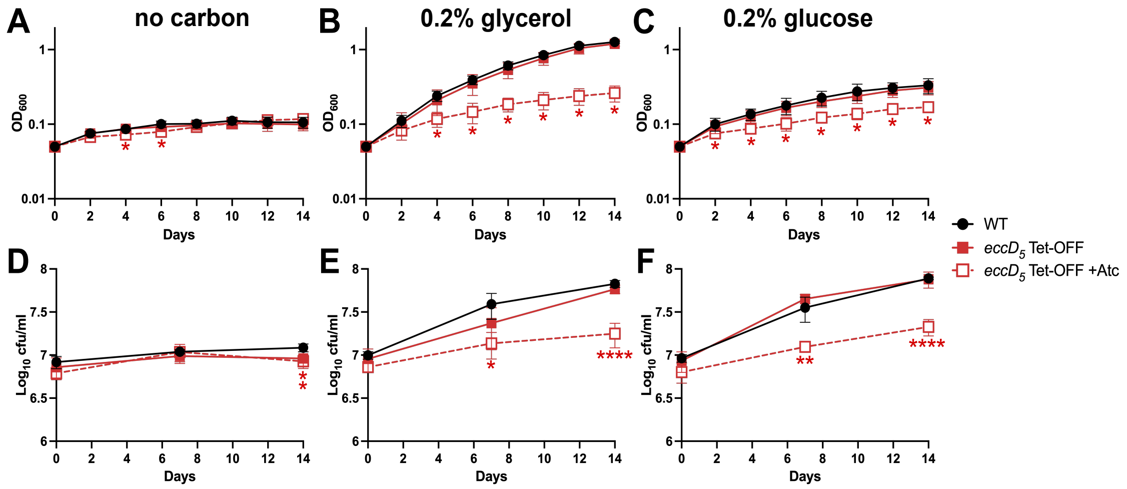


**Fig S2. *M. tuberculosis* requires ESX-5 for *in vitro* growth in minimal medium with glycerol or glucose.** WT Erdman and *eccD*$_5$ Tet-OFF were grown in complete Middlebrook 7H9 ± 100 ng/ml Atc to mid-exponential phase, then washed and diluted to $OD_{600}$ = 0.05 in minimal salts medium with 0.05% tyloxapol and no added carbon (**A, D**), 0.2% glycerol (**B, E**), or 0.2% glucose (**C, F**). Fresh Atc (100 ng/ml) was added to +Atc cultures every 7 days. Growth was monitored by $OD_{600}$ measurements every 2 days (**A-C**) or by plating serially diluted cultures on Middlebrook 7H10 agar every 7 days (**D-F**). Data are means ± standard deviations of 4 biological replicates from 2 independent experiments. Statistical analysis was performed to compare *eccD*$_5$ Tet-OFF ± Atc to WT by one-way ANOVA with Dunnett's correction (*$p$<0.05, **$p$<0.01, ****$p$<0.0001).

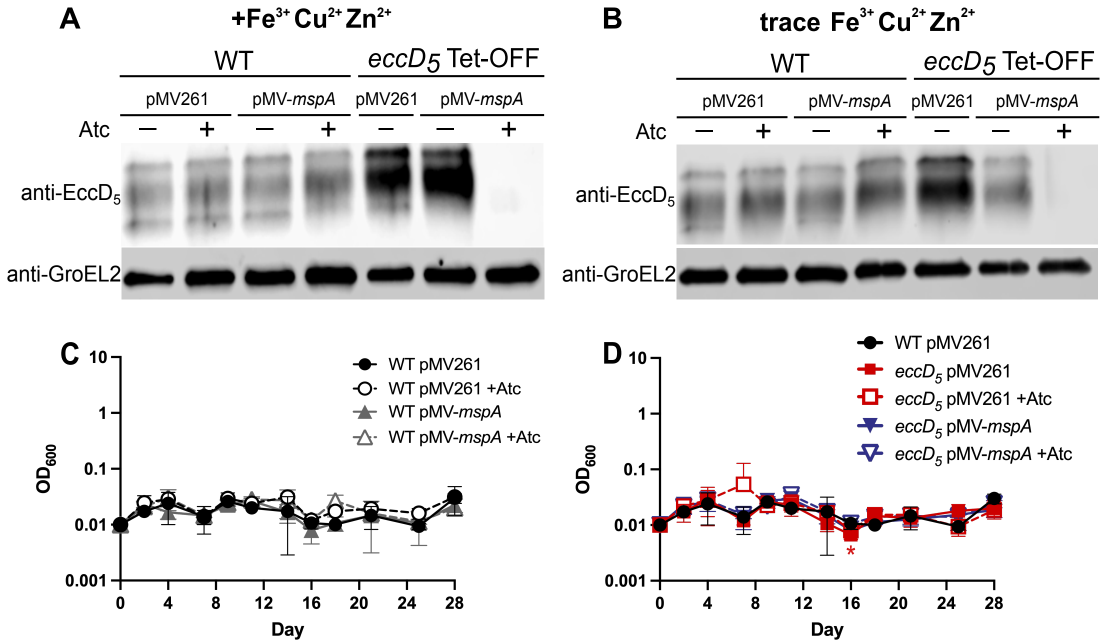


**Fig S3. Western blotting and 7H9 with no added carbon growth curve controls for *M. tuberculosis* strains expressing MspA. (A, B)** Western blots to confirm $EccD_5$ depletion. Whole cell lysates were prepared from cultures grown in 7H9 + 0.5% glycerol (**A**) or in 7H9 with trace $Fe^{3+}$, $Cu^{2+}$ and $Zn^{2+}$ + 0.5% glycerol (**B**) that are shown in Fig 2. $EccD_5$ and GroEL2 were detected in 5.8 μg total protein (**A**) or 10 μg total protein (**B**) by Western blotting. (**C,D**) The indicated strains were grown in complete Middlebrook 7H9 ± 100 ng/ml Atc to mid-exponential phase, then washed and diluted in triplicate to $OD_{600}$ = 0.01 in home-made Middlebrook 7H9 with trace $Fe^{3+}$ and $Cu^{2+}$ with 0.01% tyloxapol and no added carbon ± 100 ng/ml anhydrotetracycline (Atc). Growth was monitored by $OD_{600}$ measurements every 2-3 days. Fresh Atc (100 ng/ml) was added to +Atc cultures every 7 days. Data are means ± standard deviations. Statistical analysis was done to compare each strain and condition to the WT pMV261 untreated control (**C**) or to the *eccD5* Tet-OFF pMV261 untreated control (**D**) (*$p<0.05$; two-way ANOVA with a simple effects model and Dunnett's correction).

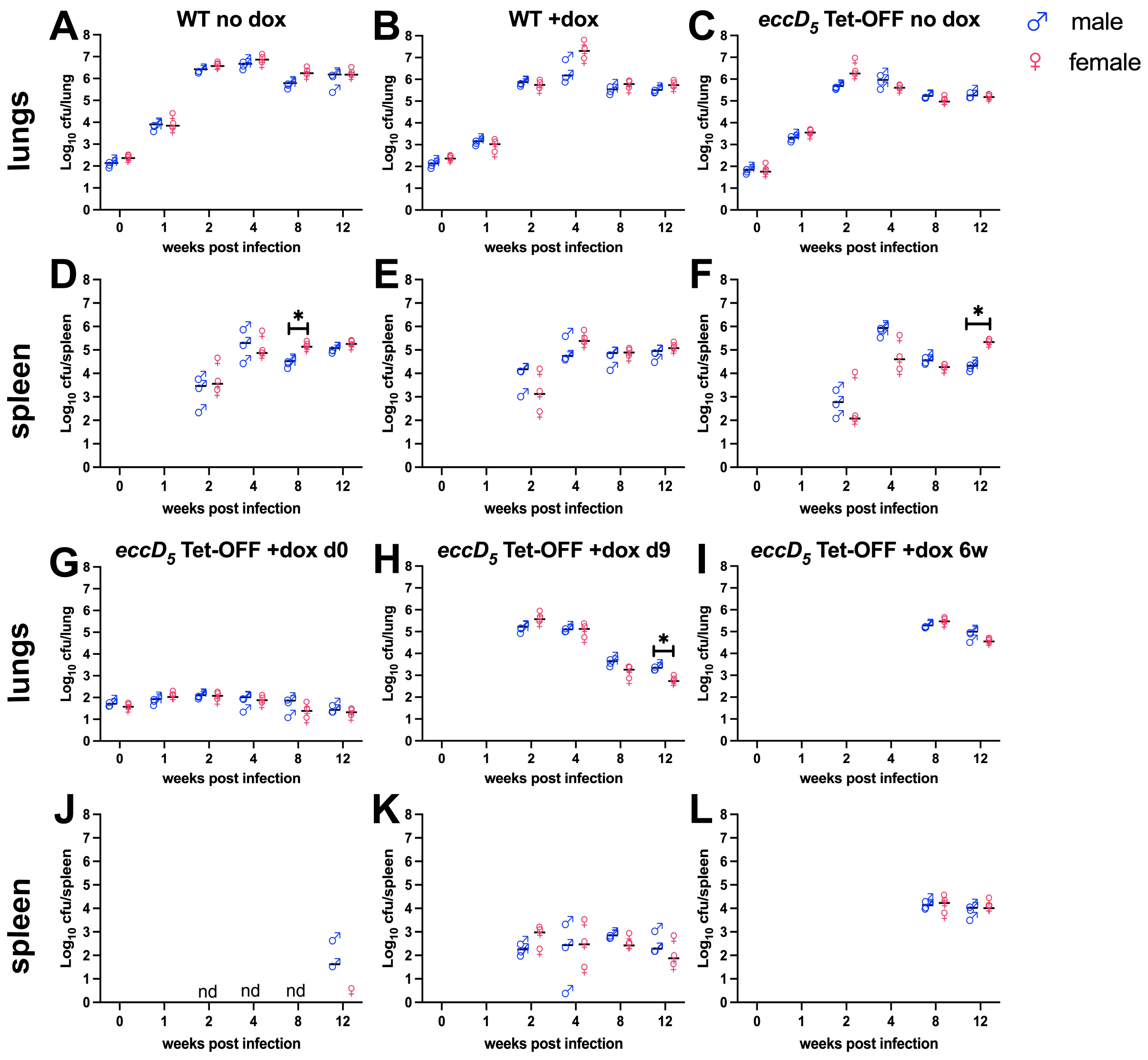


**Fig S4. *M. tuberculosis* lung and spleen burdens from individual mice, separated by sex.** Data from C57BL/6 mice that were infected by aerosol with WT Erdman (**A, B, D, E**) or *eccD*$_5$ Tet-OFF (**C, F, G-L**) and treated with doxycycline (2,000 ppm in chow) starting at the indicated time points, from Fig 5. *M. tuberculosis* CFU in lung (**A-C, G-I**) and spleen (**D-F, J-L**) tissues were determined by plating and data from male and female mice (*n*=3 each) are shown separately. Lines indicate the mean. In **J**, n.d. indicates none detected in either male or female mice (detection limit = 3 CFU). Asterisks indicate statistically significant differences between male and female mice in each experimental group (*$p$<0.05; unpaired t-test).

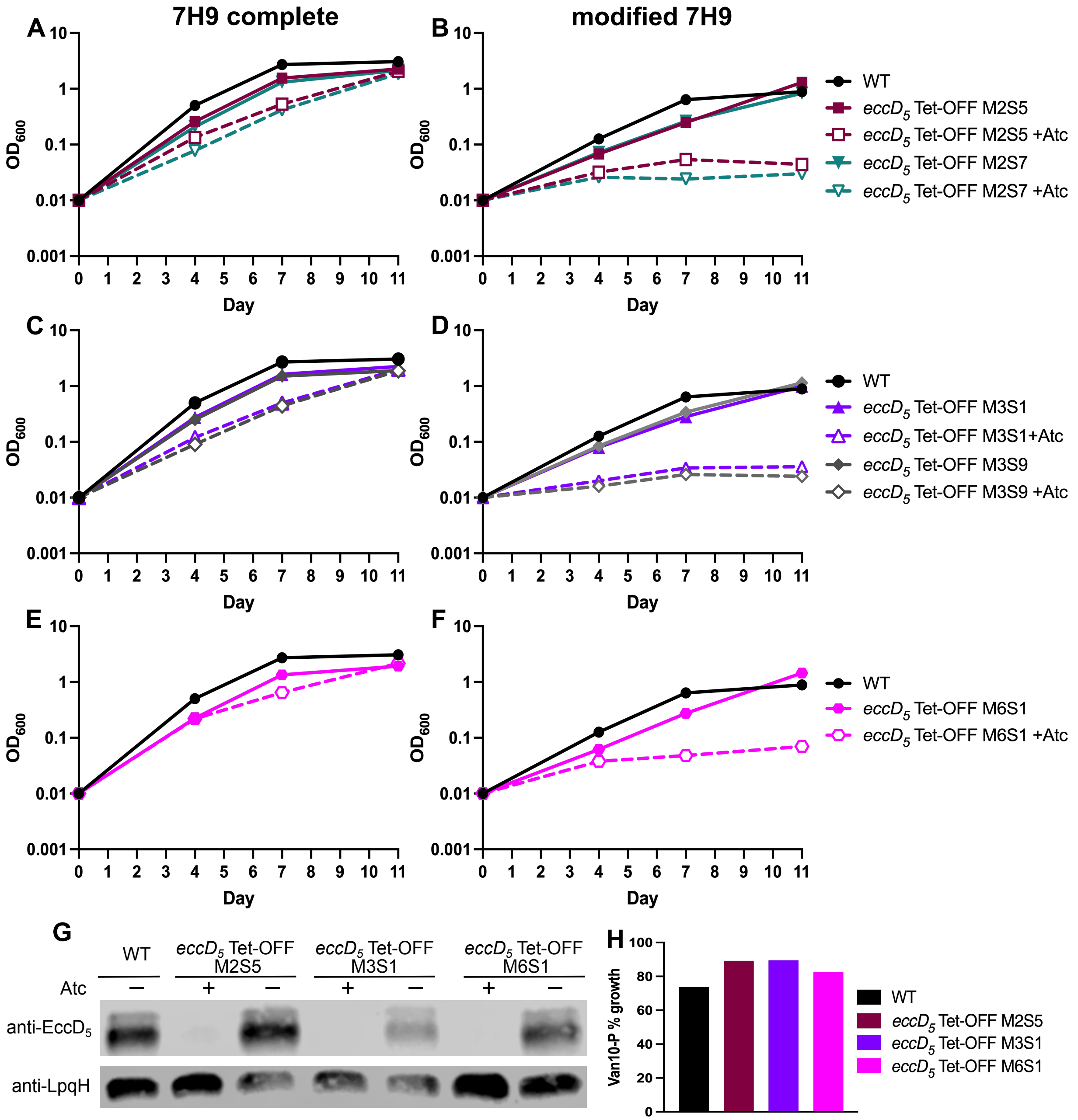


**Fig S5. *M. tuberculosis eccD$_5$* Tet-OFF isolates recovered from spleens of dox-treated mice at 12 weeks post-infection retain Atc-repressible EccD$_5$ expression.** **(A-F)** *M. tuberculosis eccD$_5$* Tet-OFF mouse spleen isolates or WT control were grown in complete Middlebrook 7H9 ± 100 ng/ml Atc to mid-exponential phase, then washed and diluted to $OD_{600}$ = 0.01 in complete 7H9 (**A, C, E**), or modified 7H9 with 0.01% tyloxapol +0.5% glycerol +0.2% glucose (**B, D, F**) ± 100 ng/ml anhydrotetracycline (Atc). Growth was monitored by $OD_{600}$ measurements every 3-4 days. Fresh Atc (100 ng/ml) was added to +Atc cultures at day 7. (**G**) *M. tuberculosis eccD$_5$* Tet-OFF mouse spleen isolates and the WT control were grown in complete 7H9 ± 100 ng/ml Atc and whole cell lysates were prepared. EccD$_5$ and LpqH were detected in 11.9 μg of total protein by Western blotting. (**H**) Vancomycin susceptibility in the presence of propionate quantified by Van10-P assay. Van10-P % growth is [($OD_{600}$ 10 μg Van)/($OD_{600}$ 0 μg Van)] x100.
